# Life histories as mosaics: plastic and genetic components differ among traits that underpin life-history strategies

**DOI:** 10.1101/2021.02.12.430905

**Authors:** Anja Felmy, David N. Reznick, Joseph Travis, Tomos Potter, Tim Coulson

## Abstract

Life-history variation reflects phenotypic variation across suites of traits. Differences among life-history strategies result from genetic differentiation, phenotypic plasticity, and genotype-by-environment interactions. If the relative strength and direction of these components differed among traits underlying a strategy, life histories might not evolve as a cohesive unit.

We tested this hypothesis on the high- and low-predation ecotypes of Trinidadian guppies, defined by distinct life-history strategies. Using common garden experiments, we assessed how strongly 36 traits were determined by ancestral habitat (i.e., ecotype) or food availability, a key environmental difference between ecotypes. Our dataset was large (1178 individuals) and included six putatively independent origins of the derived ecotype.

Traits could be confidently assigned to four groups, defined by highly significant effects of only food (13 traits), only habitat (6), both (6), or neither (11), revealing substantial variation among traits in levels of genetic and environmental control. Ecotype-food (i.e., genotype-by-environment) interactions were negligible. The directions of plastic and genetic effects were usually aligned.

This suggests that life histories are mosaics with unequal rates of phenotypic and evolutionary change. Broadly speaking of “life-history evolution” masks a complex interplay of genes and environment on the multiple traits that underpin life-history strategies.

## Introduction

Life-history strategies of populations often differ consistently between habitats. The life histories and phenotypes associated with specific habitats are known as “ecotypes”, and may evolve repeatedly in parallel. Examples include the “wave” and “crab” ecotypes of the rough periwinkle (Butlin et al. 2014), the “dwarf” and “normal” ecotypes of Lake whitefish (Campbell and Bernatchez 2004), and the “lake” and “stream” ecotypes of three-spined stickleback (Stuart et al. 2017).

Life-history differences between ecotypes can result from genetic differentiation, phenotypic plasticity, genotype-by-environment interactions, and developmental noise (Stearns 1992). Early studies of ecotypic variation often focused on demonstrating the genetic component to variation among ecotypes (e.g., Clausen et al. 1941; Reznick and Endler 1982). Later studies explored whether genetic covariances among traits constrain or facilitate the evolution of ecotypes (e.g., Karlsson Green et al. 2016; Leinonen et al. 2011), and how much phenotypic variation in key traits might be explained by salient environmental differences among habitats (e.g., Landy and Travis 2015; Trexler and Travis 1990).

When studying the evolution of ecotypes, it is important to consider all potential sources of phenotypic variation. The direction and magnitude of the components of variation have important consequences for trait evolution and population dynamics, and potentially for a species’ adaptive potential and resilience to environmental change (Coulson et al., submitted; Gienapp et al. 2008; Merilä and Hendry 2014; Robinson and Dukas 1999; West-Eberhard 2003). For example, when plasticity brings the mean phenotype closer to the optimum, as is often assumed to be the case when plastic and genetic differences are the same sign (so-called co-gradient variation), the selection differential is reduced and adaptive change proceeds more slowly (Coulson et al. 2017). By contrast, when plasticity moves the mean phenotype further from the optimum, as when plastic and genetic responses have different signs (counter-gradient variation), the selection differential is increased and adaptive change proceeds more rapidly (Coulson et al., submitted; Ghalambor et al. 2007).

The direction and strength of plastic and genetic components of life-history differences take on greater importance when considering the clusters of traits that define ecotypes. Although genetic correlations among traits can channel the direction of evolution, environmental correlations can substantially alter multivariate selection gradients (Roff 1997; Travis et al. 1999). Differences among traits within a cluster in the direction and strength of their components will influence how the cluster itself evolves – that is, through which combination of direct and indirect selection, in which order, and how quickly the traits contribute to phenotypic and genetic divergence.

The “high-predation” and “low-predation” ecotypes of Trinidadian guppies (*Poecilia reticulata*) are an example of rapid, repeatable evolutionary change in the wild (Reznick 1982; Reznick and Endler 1982; Reznick and Bryga 1996; Reznick et al. 1997). These ecotypes are adapted to different sections of natural streams in the Northern Range Mountains of Trinidad. In downstream river sections below barrier waterfalls, “high-predation” (HP) guppies coexist with several highly piscivorous fishes, such as pike cichlids (*Crenicichla alta*) and wolf fish (*Hoplias malabaricus*) (Gilliam et al. 1993). Further upstream, waterfalls gradually exclude large predators, and fish communities become dominated by “low-predation” (LP) guppies and killifish (*Rivulus hartii*) (Gilliam et al. 1993).

The most obvious difference between up- and downstream habitats lies therefore in the predator-induced mortality risk, but the contrast is more nuanced than that. Guppies attain far higher population densities in upstream habitats, leading to higher guppy biomass per unit volume (Reznick et al. 2001). In addition, upstream sites have lower algal standing crops (Grether et al. 2001) and a lower biomass of invertebrate prey (Zandonà et al. 2017). These differences presumably arise because of correlated variation in HP and LP communities between predation risk and the environment, with smaller streams, denser canopy cover, lower light levels, and thus lower primary productivity further upstream. Consequently, per-capita food availability is significantly lower in upstream habitats.

The HP and LP ecotypes have distinct life histories consistent with selection for either fast or slow reproduction, respectively (Reznick 1982; Reznick and Bryga 1996). HP guppies tend to have younger ages and smaller sizes at maturity, produce more, but smaller, offspring per litter, have shorter inter-birth intervals, and invest more resources in reproduction than LP guppies (Reznick and Endler 1982; Reznick and Bryga 1987; Reznick et al. 1990; Reznick et al. 1996). Recent work suggests that the repeated evolution of the LP life-history strategy from HP ancestors occurs primarily in response to increased population density and subsequent decreases in food availability, rather than as a direct effect of reduced predation risk (Bassar et al. 2013; Reznick et al. 2019; Reznick and Travis 2019).

Increased population density and the resulting change in food availability could influence the expression of phenotypic traits in three separate, non-exclusive ways (Auer et al. 2020). First, by reducing per-capita food availability, they may lead to plastic changes in resource-dependent traits. Second, by selecting for phenotypes optimally adapted to low-resource, high-competition environments, they may cause genetic changes. Third, genotypes may respond differently to increased population density, generating different phenotypes via genotype-by-environment interactions. If the relative strength and direction of plastic and genetic changes, or the amount of additive genetic variance, were to differ among traits that comprise the phenotype, life-history strategies may actually be mosaics of traits with unequal rates of evolution.

Here we report a test of this idea using the divergent life-history strategies of HP and LP guppies. For each of 36 traits underlying life-history variation in guppies, we evaluated the extent to which differences between guppy ecotypes are due to phenotypic plasticity in response to food level, genetic differences between fish from contrasting habitats, or genotype-by-food-level interactions. We then compared the relative directions of genetic and plastic changes, to assess whether plasticity was more likely to facilitate or constrain evolutionary change in each trait. As traits may be non-independent, we also estimated their phenotypic correlation matrix. Our hypotheses were as follows:

(H1) Few traits, if any, will be unaffected by food levels.

(H2) Size-related traits will depend on both food availability and ancestral habitat.

(H3) Where both plastic and genetic effects exist, their directions will usually be aligned (i.e., co-gradient variation).

(H4) Effects of food levels will be stronger in HP than in LP fish (significant interaction between food availability and habitat type).

We expected food effects to be ubiquitous (H1) because life-history traits are emergent properties of how the whole phenotype interacts with the environment (Coulson et al. 2006), an important aspect of which is resource availability. This is particularly true for traits related to body size (H2), which are determined by individual growth rates, which in turn depend on the resources individuals are able to accrue (Coulson 2020). Simultaneously, body size and related morphological traits, along with behavioural and physiological traits, tend to be comparatively strongly heritable, while life-history traits tend to have lower heritabilities (Postma 2014) because of their higher sensitivity to environmental effects (Hansen et al. 2011). We expected plastic and genetic differences to be the same sign (H3) based on the assumption that the genetic divergence between guppy ecotypes was often directly caused by changes in per-capita food availability (Bassar et al. 2013; Reznick et al. 2019; Reznick and Travis 2019). Lastly, we expected LP populations to cope better with low food availability than HP populations (H4), as selection for efficient resource accrual is presumably stronger in low-than in high-resource environments.

Altogether, we found that the two life-history strategies of Trinidadian guppies emerge from traits with very different levels of genetic and environmental control, suggesting that individual traits may evolve at unequal rates and yet form the ecotypes that are so readily apparent.

## Materials and Methods

### Overview of methods

We used four datasets from four independent experiments with identical design (details in Supporting Methods 1 and Table S1), and so our dataset is unusually comprehensive (sample size: 708 females and 470 males). Datasets 1, 2 and 4 have previously been published, while dataset 3 is published here for the first time. Our combined dataset encompasses guppies from seven HP and nine LP populations situated in a total of seven drainages (see map in Fig. S1). These drainages represent at least six putatively independent evolutionary origins of the LP ecotype, giving us ample power to assess which life-history traits show patterns with respect to food and habitat that are general across drainages (i.e., parallel evolution), or unique to individual drainages (i.e., genetic divergence).

Each dataset consists of a factorial experiment in the laboratory, in which two levels of a daily food ration (high vs. low) were crossed with two ancestral habitats (high-vs. low-predation). Males were measured for five, and females for 31 life-history traits. Details of the laboratory rearing protocol and of the measurement of traits can be found in the Supporting Methods 2 and 3. By using second-generation laboratory-reared fish derived from field-caught females, keeping fish in a common environment on controlled amounts of food, and splitting pairs of full-siblings between food levels, we eliminated maternal, environmental, and other non-heritable sources of variation to the greatest extent. Any trait differences between fish from HP and LP habitats therefore indicate genetic differences between ecotypes. Effects of food levels are evidence of phenotypic plasticity, and differences between ecotypes in their response to food levels imply genotype-by-food-level interactions.

### Statistical analysis

#### Overview of analyses

We conducted analyses of datasets on their own and in combination. Because datasets used different daily rations of food, we first analysed growth rates to assess the degree of differences in food levels among datasets (details in Supporting Methods 4). We then tested for effects of food availability and ancestral habitat on life-history traits by fitting linear mixed-effects models for each trait. Finally, we assessed the phenotypic correlation structure among traits to explore patterns of non-independence.

We considered an effect as highly significant only when its *p*-value was < 0.000001, to control for an elevated rate of type I errors due to running the same test on 36 traits. The resulting division of traits into those with highly significant effects of food levels and/or ancestral habitats, and those with less- or non-significant effects, was used for clustering purposes only. It is not meant to downplay weaker effects, but rather to highlight the variation among traits in the strength of their dependence on the two predictors. In all linear models we looked for outliers using function “CookD” in R-package “predictmeans” (Luo et al. 2018) and by comparing models without potential outliers to models including them.

However, no data point lay outside 0.5 Cook’s distance (maximum Cook’s distance: 0.20) and excluding the few less-extreme outliers that were present did not change our results. We thus retained all data in our final analyses.

Statistical analyses were performed in R v. 3.4.0 (R Core Team 2017) and 4.0.0 (R Core Team 2020). Scatter plots were prepared using R-package “beeswarm” (Eklund 2016), the heat map using R-packages “gplots” (Warnes et al. 2019) and “RColorBrewer” (Neuwirth 2014), and network plots using R-package “igraph” (Csardi and Nepusz 2006).

### Effects of food level and ancestral habitat

When analysing the datasets jointly, we fitted a separate linear mixed-effects model for each of the 36 traits to investigate effects of experimental food levels and ancestral habitats, while controlling for differences between datasets, drainages, and maternal identities. Table 1 provides information on the traits that were studied, and on how they were modelled. Details of the model selection process can be found in the Supporting Methods 5. All models were fitted using R-packages “lme4” (Bates et al. 2015) and “lmerTest” (Kuznetsova et al. 2017). We tested the significance of random effects by means of log-likelihood ratio tests comparing the full model to one without the random effect in question. After selecting the best-fitting model for each measured trait, we evaluated the plausibility of the model results by comparing them to plots of trait values against all predictors, separately for models with and without two-way interactions.

**Table 1.**
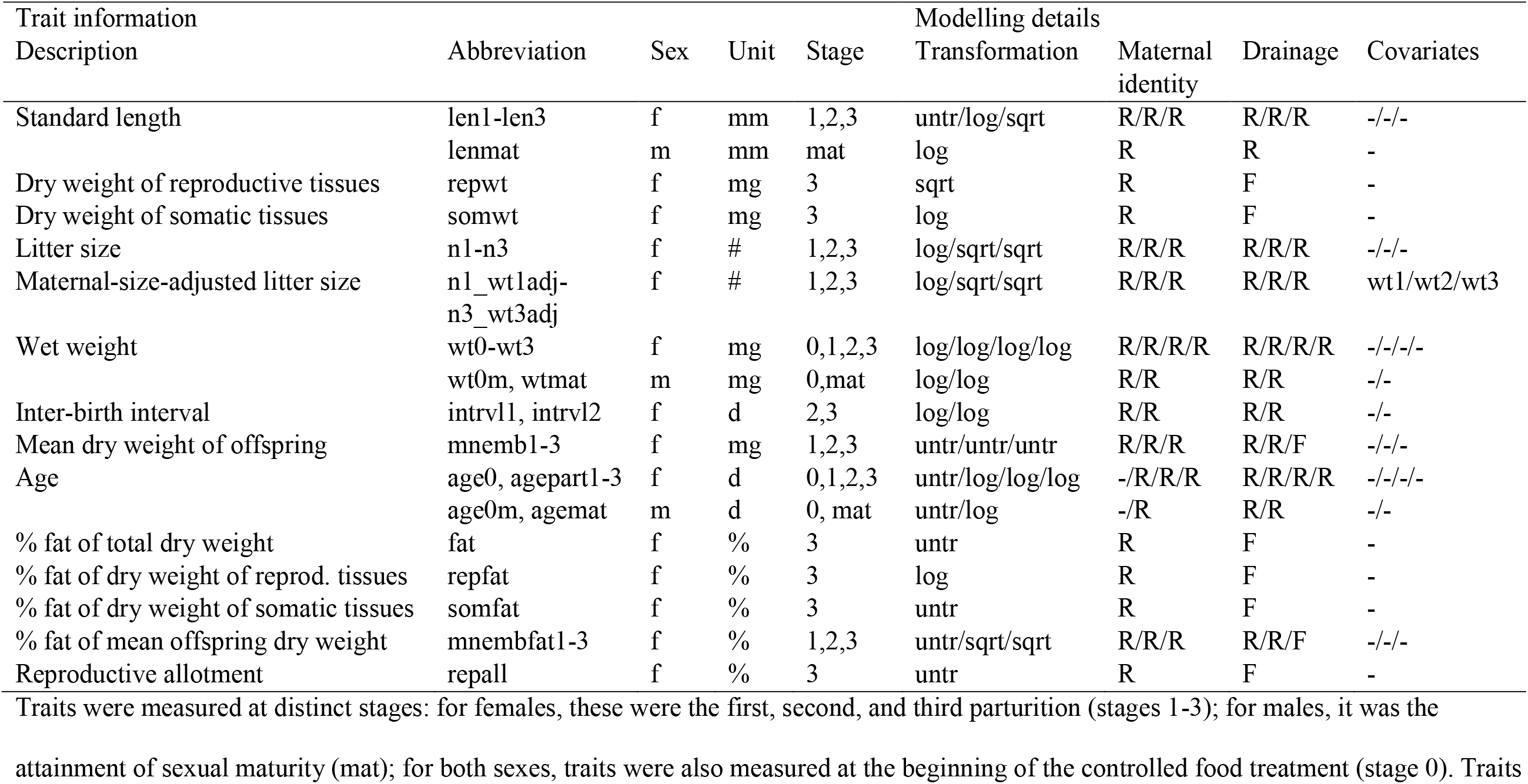
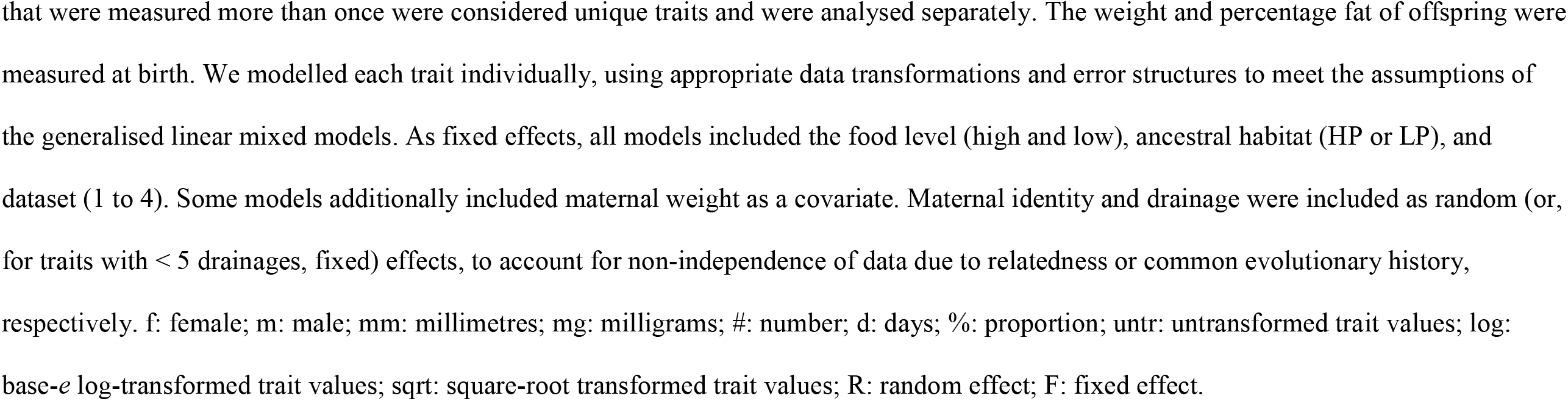
Traits considered in this study, and details of how they were modelled.

| Trait information |  |  |  |  | Modelling details |  |  |  |
| --- | --- | --- | --- | --- | --- | --- | --- | --- |
| Description | Abbreviation | Sex | Unit | Stage | Transformation | Maternal identity | Drainage | Covariates |
| Standard length | len1-len3 | f | mm | 1,2,3 | untr/log/sqrt | R/R/R | R/R/R | -/-/- |
|  | lenmat | m | mm | mat | log | R | R | - |
| Dry weight of reproductive tissues | repwt | f | mg | 3 | sqrt | R | F | - |
| Dry weight of somatic tissues | somwt | f | mg | 3 | log | R | F | - |
| Litter size | n1-n3 | f | # | 1,2,3 | log/sqrt/sqrt | R/R/R | R/R/R | -/-/- |
| Maternal-size-adjusted litter size | n1_wt1adj-<br>n3_wt3adj | f | # | 1,2,3 | log/sqrt/sqrt | R/R/R | R/R/R | wt1/wt2/wt3 |
| Wet weight | wt0-wt3 | f | mg | 0,1,2,3 | log/log/log/log | R/R/R/R | R/R/R/R | -/-/-/- |
|  | wt0m, wtmat | m | mg | 0,mat | log/log | R/R | R/R | -/- |
| Inter-birth interval | intrvl1, intrvl2 | f | d | 2,3 | log/log | R/R | R/R | -/- |
| Mean dry weight of offspring | mnemb1-3 | f | mg | 1,2,3 | untr/untr/untr | R/R/R | R/R/F | -/-/- |
| Age | age0, agepart1-3 | f | d | 0,1,2,3 | untr/log/log/log | -/R/R/R | R/R/R/R | -/-/-/- |
|  | age0m, agemat | m | d | 0, mat | untr/log | -/R | R/R | -/- |
| % fat of total dry weight | fat | f | % | 3 | untr | R | F | - |
| % fat of dry weight of reprod. tissues | repfat | f | % | 3 | log | R | F | - |
| % fat of dry weight of somatic tissues | somfat | f | % | 3 | untr | R | F | - |
| % fat of mean offspring dry weight | mnembfat1-3 | f | % | 1,2,3 | untr/sqrt/sqrt | R/R/R | R/R/F | -/-/- |
| Reproductive allotment | repall | f | % | 3 | untr | R | F | - |

Traits were measured at distinct stages: for females, these were the first, second, and third parturition (stages 1-3); for males, it was the
attainment of sexual maturity (mat); for both sexes, traits were also measured at the beginning of the controlled food treatment (stage 0). Traits
that were measured more than once were considered unique traits and were analysed separately. The weight and percentage fat of offspring were measured at birth. We modelled each trait individually, using appropriate data transformations and error structures to meet the assumptions of the generalised linear mixed models. As fixed effects, all models included the food level (high and low), ancestral habitat (HP or LP), and dataset (1 to 4). Some models additionally included maternal weight as a covariate. Maternal identity and drainage were included as random (or, for traits with < 5 drainages, fixed) effects, to account for non-independence of data due to relatedness or common evolutionary history, respectively. f: female; m: male; mm: millimetres; mg: milligrams; #: number; d: days; %: proportion; untr: untransformed trait values; log: base-*e* log-transformed trait values; sqrt: square-root transformed trait values; R: random effect; F: fixed effect.

For each trait, we obtained mean-standardised effect sizes for experimental food levels and types of ancestral habitat from the best-fitting model. These were computed by dividing the model estimate (i.e., the partial regression coefficient) for the effect of food or habitat, respectively, by the intercept, and multiplying this ratio by 100.

In addition to analyses using the combined datasets, we conducted dataset-specific analyses. For each dataset and trait present within the dataset, a separate model was fitted, resulting in a total of 112 models (details in Supporting Methods 6).

### Phenotypic correlations

We assessed the phenotypic correlation structure between traits by computing Pearson product-moment correlation coefficients between all pairs of traits, estimating correlations separately for traits measured in males and females. Correlations shown in the main text were calculated using all our data, but correlations including only fish from a given food level and habitat type can be found in the Supporting Information.

## Results

### Summary of key results

Life-history traits could be assigned to one of four groups, characterised by highly significant effects of food alone (13 traits), ancestral habitats alone (6 traits), both food and habitats (6 traits), and neither (11 traits). Size-related traits primarily depended on food, inter-birth intervals and offspring size primarily on habitat, and reproductive timing on both. Litter size depended on food, and, when adjusted for maternal size, also on habitat. The classification of traits into groups based on which factors were important was reinforced by their phenotypic correlation structure: traits were strongly correlated with each other within but not between groups. Where both plastic and genetic changes existed, they pointed in the same direction, with few exceptions. Interactions between food levels and habitat types were either absent (31 traits) or only weakly significant (5 traits). Trait differences between datasets reflected variation in experimental food levels. There was significant variation among drainages in both mean trait values and the overall patterns with respect to food levels and habitat types.

### Differences in food levels between datasets

Absolute food levels were not consistently repeated across datasets 1 to 4, yet the observed patterns of female growth between the start of experimental food treatments and the birth of litter 3 were repeatable (Fig. S2, Supporting Results, Tables S2 and S3).

### Effects of food level and ancestral habitat

Twenty-five out of 36 phenotypic traits (69.4%) showed highly significant effects of either the experimental food treatment, the ancestral habitat, or both (Fig. 1, full model results in Tables S4-S39). We could classify traits into four groups, based on the statistical significance of food and habitat effects, with classifications reflected in the sizes of effects. Group A traits were heavily affected by food but not the ancestral habitat (13 traits). For example, fish were, on average, 4.2% shorter and 5.2% lighter at the lower food level, while they were only 0.6% longer and 1.3% heavier when from LP rather than HP habitats (mean-standardised effect sizes; Fig. 1). Group B traits were heavily affected by the habitat but not food (6 traits): In fish from LP habitats, inter-birth intervals were 2.7% longer and new-born offspring 26.1% heavier than in fish from HP habitats, compared to food-level differences of only 1.1% and 6.9%, respectively. Group C traits showed strong effects of both food and habitat (6 traits), with, for instance, fish maturing 2.7% more slowly at the lower food level and 2.3% more slowly when from LP habitats. In group D, traits lacked highly significant effects of either food levels or habitat types (11 traits). Representative examples of traits (Fig. 2a-d) illustrate how group A traits showed near-identical mean values in HP and LP fish but a clear difference between food levels, while group B traits displayed the opposite pattern. In group C, trait means differed between all four experimental categories; in group D, all trait means were similar. The fact that 13 out of 32 traits measured after feeding regimes had begun (40.6%) were comparatively food-independent means that H1 must be rejected.

**Figure 1.**
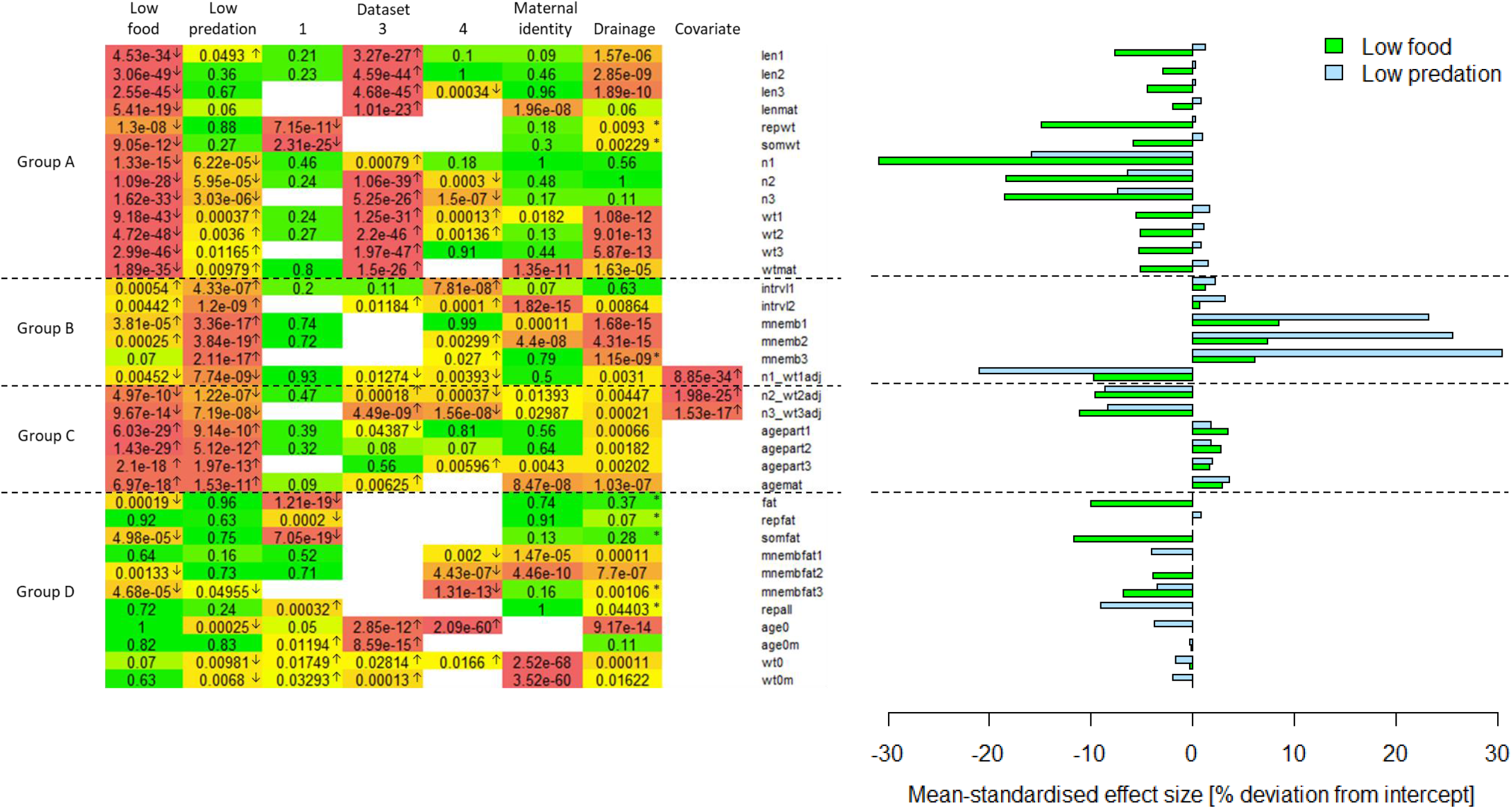
Heat map of *p*-values (left) and effect sizes (right) of experimental food levels and ancestral habitats on life-history traits. A separate model was fitted for each of 36 traits. Models included the experimental food level (high vs. low), the ancestral habitat (high-vs. low-predation) and the dataset (1 to 4) as categorical fixed effects, with a reference level of high food, under high predation, using dataset 2. Models of size-adjusted litter size include postpartum maternal weight as a covariate. Random effects were maternal identity nested within drainage, unless there were only four drainages, in which case drainage was fitted as a categorical fixed effect. For these traits, the *p*-value of the overall effect of drainage comes from a log-likelihood ratio tests comparing the full model to one without drainage (indicated by an asterisk). In dataset 1 fish originated from a single drainage not sampled in any other dataset, so effects of dataset 1 and drainage are confounded. For full model results, see Tables S4-S39. In the heat map, arrows indicate the direction of effects with *p* < 0.05. Traits were sorted by group, with group A = highly significant effect of only food, group B = highly significant effect of only habitat, group C= highly significant effects of both, group D = highly significant effects of neither. Effects were considered highly significant when *p* < 0.000001, to account for an increased type I error rate due to multiple testing. Note that four traits in group D (age0, age0m, wt0, wt0m) were measured before controlled food treatments began and thus serve as negative controls for food effects. Effect sizes correspond to the *p*-values for food levels and ancestral habitats shown on the heat map. For traits that were transformed before models were fitted, results are provided on the transformed scale. Trait abbreviations are explained in Table 1.

**Figure 2.**
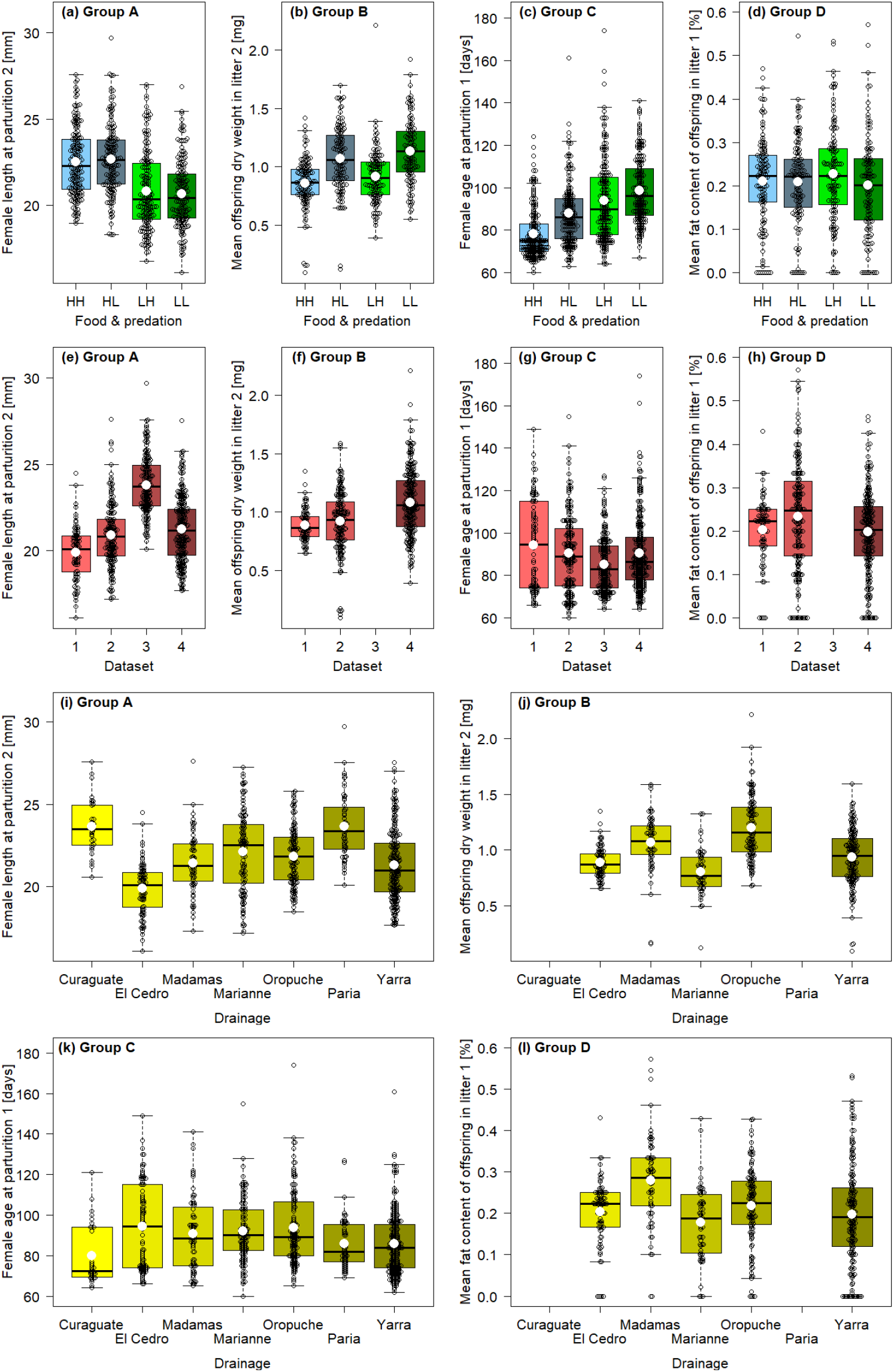
Effects of food level and ancestral habitat (a-d), dataset (e-h) and drainage (i-l) on one trait each from groups A to D. Traits were chosen because of their differential dependence on food levels and ancestral habitats, with traits being either highly significantly affected by food only (group A), habitat only (group B), both (group C), or neither (group D). White circles denote mean values of categorical predictor levels. HH: high food, high predation; HL: high food, low predation; LH: low food, high predation; LL: low food, low predation.

Families of similar traits fell into the same group (Fig. 1). For example, all length and weight measurements of adult fish were group A traits (Fig. 3), leading to the rejection of H2. Litter size forms a partial exception to this rule. Raw litter sizes belonged to group A, with strong food effects and modest but sub-threshold effects of ancestral habitats. However, litter size strongly increased with maternal body weight, with first litters containing 3.2 times more offspring in heavy females (> 250 mg) than in light females (< 150 mg; averages based on raw data). Once maternal weight was included as a covariate, habitat effects became significant and litter sizes were re-classified as group B (litter size 1) and group C traits (litter sizes 2 and 3).

**Figure 3.**
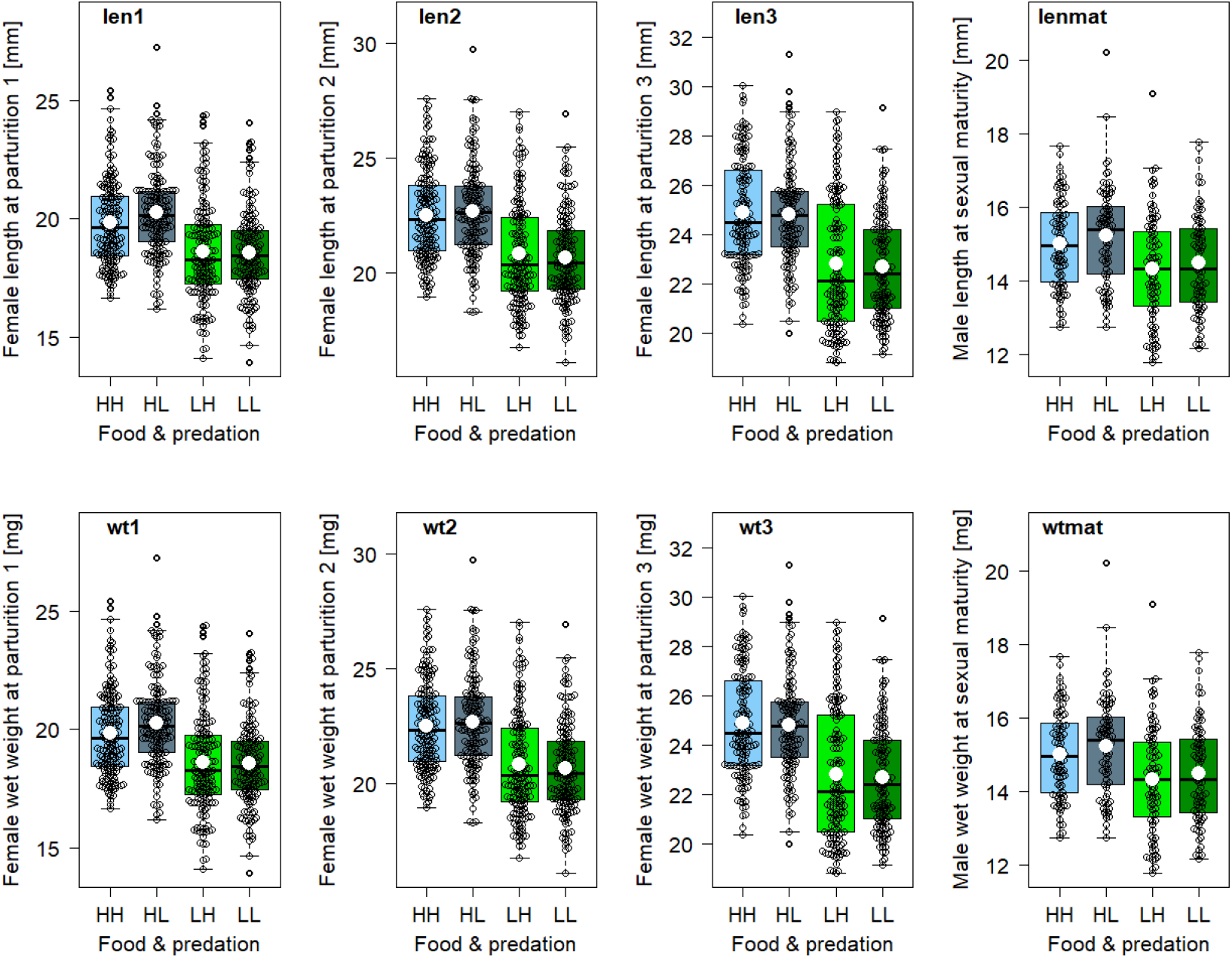
Effects of food levels and ancestral habitat on size-related traits in adult guppies. These are all group A traits, with a highly significant effect of food yet no or but weakly significant effects of habitats. White circles denote mean values of categorical predictor levels. HH: high food, high predation; HL: high food, low predation; LH: low food, high predation; LL: low food, low predation. Trait abbreviations are explained in Table 1.

The consistency of food and habitat effects on trait families is also visible in the direction of effects (see arrows and effect sizes in Fig. 1). Adult length and weight, as well as litter size, invariably decreased under low-food conditions. Inter-birth intervals were consistently longer and offspring sizes at birth larger in LP fish. All ages at reproductive events increased in response to low-food and were later in fish from LP conditions. Litter size, adjusted for maternal weight, was always smaller in LP fish. For traits with both plastic and genetic effects (group C), the directions of these effects were aligned. This is consistent with H3. Only adult wet weights showed plastic and (weakly significant) genetic responses that acted in opposition.

Dataset-specific analyses for each trait showed that these patterns were consistent across datasets. Although there were small differences in effect sizes for a few traits, effects found in the combined dataset were mirrored in individual datasets (data not shown).

### Interactions between food level and habitat type

The interaction between food treatment and ancestral habitat was absent (31 traits) or only weakly significant (5 traits, all *p* ≥ 0.0026; results of models including two-way interactions in Fig. S3). Non-significant food-habitat interactions were also found in our analyses of female growth rates (*F*_1_ ≤ 2.60, *p* ≥ 0.11; Supporting Results, Tables S2 and S3). These results remained unchanged when fitting habitat as the first predictor in analyses of variance (*F*_1_ ≤ 3.1, *p* ≥ 0.08). Accordingly, H4 must be rejected.

### Differences between datasets, drainages, and mothers

A look at the interactions between datasets and food levels, and between datasets and ancestral habitats, revealed that most responses were similar across datasets (Fig. S3). This was despite considerable differences among datasets in mean trait values, which were most notable in the strongly food-dependent group A traits (Figs. 1, 2e-h). For example, in the unpublished dataset 3, where food levels were highest, adult fish were significantly larger and offspring numbers higher than in dataset 2 (the reference level). It should be noted that datasets also differed in the drainages and localities that were sampled; however, as our models included drainage as an additional predictor (see below), we are confident that the dataset effects found here mostly reflect variation in food levels.

The drainage of origin of experimental fish significantly affected some traits, particularly in groups A and B (Fig. 1, 2i-l). For instance, at parturition 1, females from the Yarra drainage were 6.3% smaller, 22.9% lighter, and produced embryos that weighed 20.9% less, than females from the Oropuche drainage (averages based on raw data), although these fish were all part of dataset 4 and hence did not differ in food levels. Importantly, there was also variation among drainages in the overall patterns with respect to food and ancestral habitat (see Figs. S4-S10 for seven traits showing drainage-specific effects of varying strength). For example, male length at sexual maturity was larger in LP than in HP guppies from drainages Madamas and Yarra (dataset 2), yet smaller in LP than in HP guppies from drainage Marianne (dataset 2), and identical in LP and HP guppies in another locality sampled within drainage Marianne (unpublished dataset 3, Fig. S6).

Maternal identity explained part of the variation in male length, weight and age at sexual maturity, and most significantly in the weight of both males and females before the onset of experimental treatments (Fig. 1). The influence of maternal identity on other trait families was mixed, with some traits affected by it (e.g. inter-birth interval 2) but closely related ones not (e.g. inter-birth interval 1). Note that maternal identity is confounded with variation in the microenvironment fish experienced in the laboratory.

### Phenotypic correlation structure

All except nine traits showed moderate to strong phenotypic correlations to other traits (correlation coefficient *r* ≥ 0.5; Fig. 4). Correlations were all highly significant (*p* << 0.000001) and positive, and remained largely unchanged when calculated within a given food level and habitat type (Figs. S11-14). The correlation structure reflected the classification of traits into the aforementioned, statistical groups A-D: strong correlations were only present between traits belonging to the same group (see colours in Fig. 4). Particularly groups A and C contained tight clusters of inter-correlated traits. In group B correlations were weaker and scarcer, but still present between individual pairs of traits. In contrast, most group D traits, such as the percentage fat of offspring, the reproductive allotment, and the weight at the beginning of the food treatment, lacked correlations to other traits.

**Figure 4.**
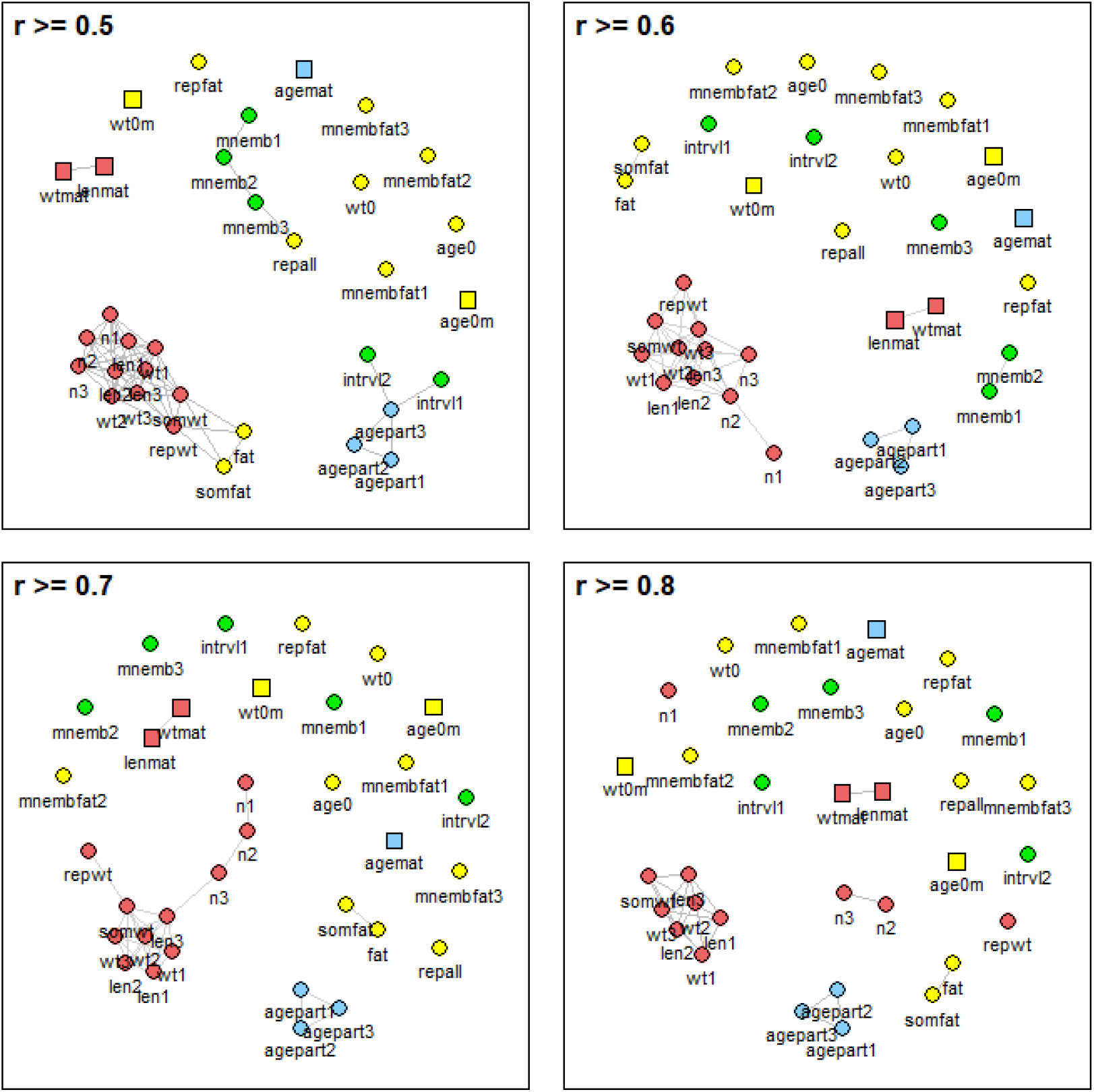
Network plots of phenotypic correlations between life-history traits. Grey lines connect traits with pairwise Pearson product-moment correlation coefficients of *r* ≥ 0.5, ≥ 0.6, ≥ 0.7 or ≥ 0.8. No lines were plotted for correlation coefficients of *r* < 0.5. The correlation between traits measured in females (circles) and males (squares) is *r* = 0 by definition. The position of traits and trait clusters relative to one another is irrelevant. Colours mark groups of traits with similar statistical dependence on experimental food levels and ancestral habitats, with group A (red) = highly significant effect of food only, group B (green) = highly significant effect of habitat only, group C (blue) = highly significant effect of both, group D (yellow) = highly significant effect of neither (see Fig. 1). Trait abbreviations are explained in Table 1.

## Discussion

The different life-history strategies of Trinidadian guppies inhabiting high- and low-predation habitats are well-documented. Here we used this system to examine whether the individual traits that make up a life-history strategy have similar levels of genetic and environmental control. We found that life-history traits could be assigned to four groups, defined by highly significant effects of only food, only habitat, both, or neither, showing that the relative contribution of genetic versus environmental components to the divergence of ecotypes varied substantially among traits.

The classification into four groups suggested itself because of the clear distinction between highly significant effects on some traits, and weakly or non-significant effects on others. Group assignments were corroborated by strong phenotypic correlations of traits within but not between groups, and were largely consistent across datasets despite differences in food levels. Families of related traits pertained to the same group and showed effects that pointed in the same direction. This was true not only for repeated measures that inevitably increased over time (e.g., female weight at parturition 1-3), but for mathematically independent traits as well (e.g. offspring weights in litters 1-3). Male and female counterparts of traits fell into the same group, despite, by design, non-existent phenotypic correlations between traits measured in separate sexes.

### Life histories as mosaics with potentially unequal rates of evolution

The variation in the magnitude and direction of the traits’ plastic and genetic components almost certainly has consequences for life-history evolution. Phenotypic change will be fast if a trait is predominantly plastic or if plastic and genetic responses are aligned (co-gradient variation; Conover and Schultz 1995; Levis and Pfennig 2016; Robinson and Dukas 1999; West-Eberhard 2003). Genetic change, on the other hand, will be fast if plastic and genetic responses have different signs (counter-gradient variation), and medium-fast if they are the same sign, but plasticity uncovers cryptic genetic variation via strong genotype-by-environment interactions (Coulson et al. 2017; Coulson et al., submitted; Ghalambor et al. 2007).

Our results suggest that life-history strategies in Trinidadian guppies are mosaics of traits with different rates of phenotypic and genetic change. As HP guppies colonise a new LP habitat, phenotypic differences to the ancestral population may develop most readily in strongly food-dependent traits: in adult body size, in litter size, and in reproductive timing. These phenotypic changes may persist, except for those in adult wet weight, the only trait showing evidence of counter-gradient variation. In adult wet weight, the plastic response may soon be offset by opposing genetic change as selection drives the mean phenotype back to a local optimum (Coulson et al. 2017). We thus tentatively predict that adult wet weight may evolve quite quickly. This is in contrast to traits that showed co-gradient variation (e.g., reproductive timing, maternal-size-adjusted litter size), where evolution may proceed rather slowly due to the lack of strong genotype-by-environment interactions. In these traits, plasticity will bring phenotypes closer to local optima for all genotypes equally, thereby masking advantageous genotypes from selection (Coulson et al. 2017). Also traits where genetic effects were weak, such as adult lengths and dry weights, are predicted to evolve slowly. Where the divergence between ecotypes was found to be primarily genetic, as for inter-birth intervals and offspring weights, the rate of evolution is expected to be intermediate, and identical to the rate of phenotypic change. Finally, neither phenotypic nor evolutionary change is expected to happen quickly in traits without either food or habitat effects.

At this time, these predictions must be considered speculative. For one thing, they assume that food availability is the most important environmental driver of life-history differences between guppy ecotypes in the field. In truth, life-history traits could be plastic with respect to other aspects of the environment, such as social density, predation risk, or parasite pressure, even though food availability is likely to be a key determinant of life histories in Trinidadian guppies (Bassar et al. 2013; Reznick et al. 2019). For another thing, our predictions assume that the magnitude of differences in laboratory food levels is comparable to or at least as great as natural variation in per-capita food levels. While our results were consistent across four datasets with unequal food levels, demonstrating their robustness to variation in food quantity, the diets of laboratory fish differed from those of free-living fish: We fed our fish high-protein liver paste and brine shrimp nauplii, rather than the low-calorie algae typically present at LP sites (Grether et al. 2001; Zandonà et al. 2017). If LP fish were better at scraping off and metabolising low-quality periphyton, our study would be underestimating the effects of differences in food levels. Neither did we measure competitive ability. By housing fish individually, we tested whether LP fish can better tolerate a low-quantity diet, which they could not, given the general lack of strong interactions between food levels and ancestral habitats. Yet, LP fish may be adapted to food scarcity in ways we did not measure, such as by feeding more quickly or being superior competitors to HP fish.

### Parallelism versus divergence in life-history evolution

For some traits, patterns of food and habitat dependence were consistent across the seven drainages included in our study, while for others they were not. Where patterns were consistent, they provide strong evidence for parallel evolution (see also Reznick and Bryga 1996; Reznick et al. 1996). However, many traits showed differences among drainages: in mean trait values, in the amount of plasticity, and, interestingly, in the magnitude of habitat (i.e., genetic) effects. In fact, variation among drainages in relative trait differences between HP and LP localities was so substantial that habitat effects frequently changed sign between drainages (Figs. S4-S10).

Spatial variation in the strength of ecotypic divergence could have multiple causes. First, drainages could vary in the history of their populations, with LP populations in some drainages being more recently derived from HP populations than others. Such differences should show up as variation among drainages in the amount of within-stream migration and in estimated divergence times between up- and downstream populations. Genetic analyses have confirmed these a posteriori hypotheses (Blondel et al. 2019; Fraser et al. 2015; Willing et al. 2010).

Second, selection pressures may differ between drainages in response to more fine-grained environmental heterogeneity. In Trinidad, canopy openness and algal standing crops differ not only between the upstream and downstream sites of a given drainage, but also between drainages (Grether et al. 2001; Reznick et al. 2001). Accordingly, selection gradients may differ among drainages, uniquely reflecting the local resource landscape.

Third, drainages may have different genetic routes via which adaptation occurs. Genomic analyses showed that guppies in different drainages are highly divergent from one another (Willing et al. 2010), suggesting that the sort of genetic variation selection can act upon might differ among drainages. Let us consider adult body size, which, depending on the drainage, was larger, smaller, or identical in HP and LP fish. Male size at maturity and female size at first parturition are determined by the intersection of growth rates with threshold rules for maturation. Threshold rules define the age and size at reproductive events (Fig. 5a, b).

**Figure 5.**
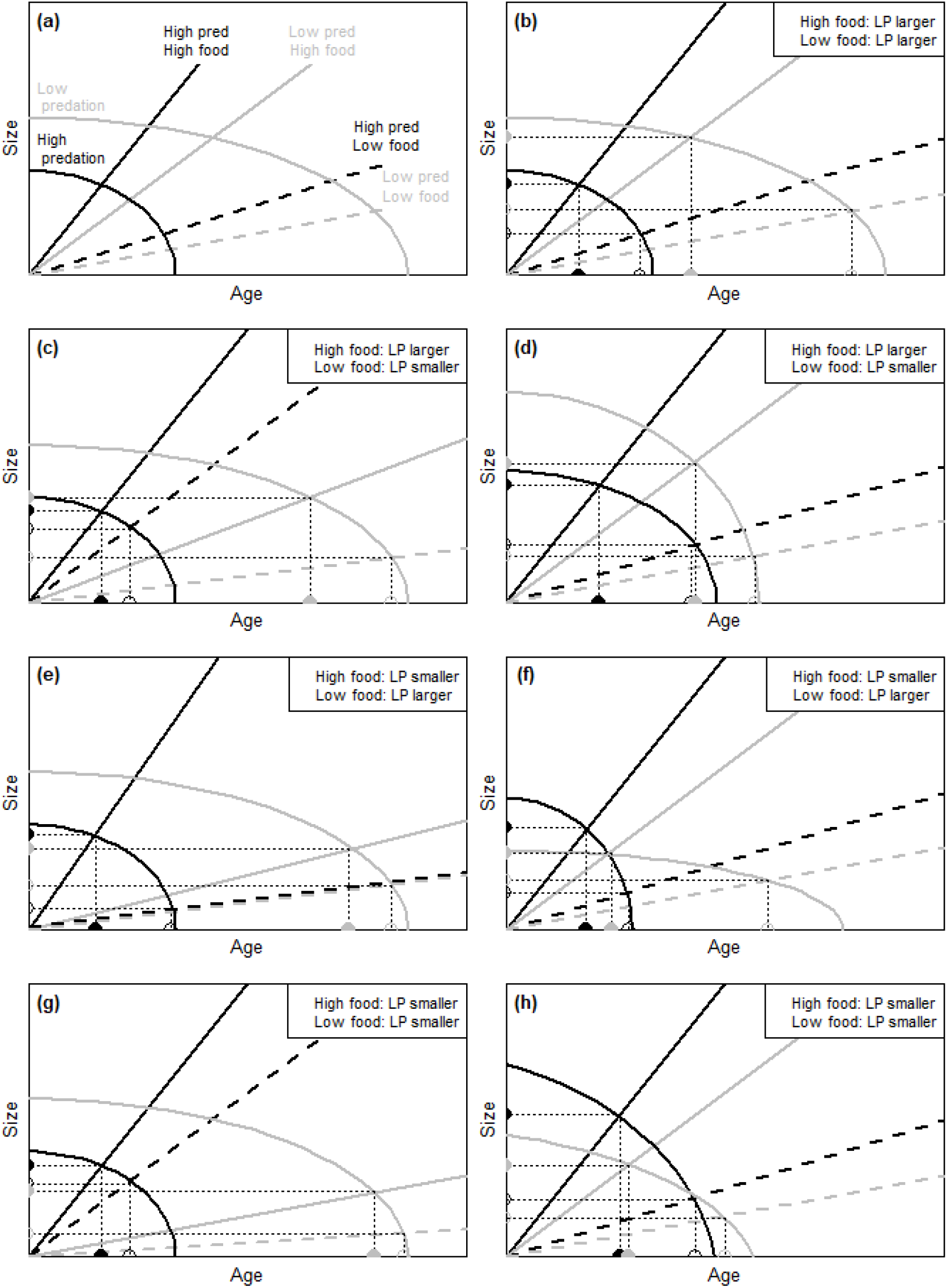
Theoretical prediction of changes in body size as a consequence of variation in growth rates (a, c, e, g) and threshold rules for maturation (b, d, f, h). Male size at sexual maturity, and female size at first parturition, are determined by the intersection of growth rates (straight lines) with threshold rules defining the age and size at reproductive events (curved lines). Fish from HP (black lines) and LP habitats (grey lines) potentially differ in threshold rules for maturation, and in how they grow innately or in response to high (solid lines) and low food availability (dashed lines). Shown is how size at maturity/first parturition reacts to changes in growth rates while keeping threshold rules constant (a, c, e, g), and to changes in threshold rules while keeping growth rates constant (b, d, f, h). Note that change in either one alone is sufficient to produce all possible ways in which sizes can differ between HP and LP populations at high and low food availability. (a, b) Same initial situation, with size at maturity/first parturition larger in LP fish irrespective of food levels. (c, d) Size still larger in LP fish under high food, but smaller in LP fish under low food. (e, f) Reversed situation. (g, h) Size smaller in LP fish at both food levels. In all scenarios, the age at maturity/first parturition is younger in HP fish, and younger under high food, in accordance with results found here and in previous studies.

When assuming that both growth rates and threshold rules vary among drainages (shown for growth rates by Arendt and Reznick 2005; Potter et al. 2019), among HP and LP habitats, and, certainly for growth rates, in response to food availability, it becomes clear how habitat effects on adult size can vary across drainages. Every possible pattern of adult size with respect to habitat and food can be produced by varying either the growth rates (Fig. 5a, c, e, g) or the threshold rules (Fig. 5b, d, f, h), demonstrating that identical phenotypes can be produced via different genetic routes, and different phenotypes via identical routes.

### Comparison with previous work

It is of interest to compare our results for habitat type with previous studies. The ideal comparisons are those based on paired populations of HP and LP guppies from the same drainage, since there is evidence of habitat-independent differences among drainages in life histories, well-illustrated for our dataset 4 (Reznick et al. 2006; Reznick et al. 2004). In this regard, our study is limited because dataset 1 compares an introduced population with the HP source population only four years after the introduction, during which only male age and size at maturity had evolved (Reznick and Bryga 1987). The equivalent evolution of female life-history traits only began to become apparent seven years post-introduction (Reznick et al. 1997). Dataset 3 included four populations from drainages represented by only one habitat type, so they confound drainage with habitat type. Only datasets 2 and 4 consist solely of paired HP and LP populations. Together, datasets 2-4 contain paired populations from four drainages – the Madamas, Marianne and Yarra in dataset 2, the Marianne in dataset 3, and the Oropuche and Yarra in dataset 4. Comparing habitat effects between our analyses and earlier studies is thus not ideal. Nevertheless, we also analysed each dataset on its own, and found our results to be largely consistent across datasets.

A superficial discrepancy between our study and earlier work is the absence of highly significant effects of habitat type on adult size. Most previous studies, whether conducted in the field (Reznick and Endler 1982; Reznick and Bryga 1987; Reznick et al. 1990; Reznick et al. 1996; Zandona et al. 2011) or laboratory (Grether et al. 2001; Potter et al. 2020; Reznick 1982; Reznick et al. 2019; Reznick and Bryga 1987; Reznick et al. 1990; Reznick and Bryga 1996), found that adult LP fish of both sexes were significantly larger and heavier than HP fish. One source of the difference in our conclusions is our stringent criterion for significance (*p* < 0.000001). For example, our data revealed a habitat effect for female wet weight (*p* = 0.00037) that would be considered highly significant in conventional data interpretations.

Another reason for the weaker apparent habitat effects on size-related traits is the aforementioned variation among drainages in habitat effects. These inconsistencies, driven by environmental (Grether et al. 2001; Reznick et al. 2001) and genetic (Fraser et al. 2015; Willing et al. 2010) differences among drainages, could mask the overall effect of habitat when measured across multiple drainages.

A second difference between this and earlier studies pertains to the partial lack of significant habitat effects on the number of offspring per litter found here. In earlier studies, LP guppies produced smaller litters in both field and laboratory comparisons (e.g., Reznick 1982; Reznick et al. 2001; Reznick and Bryga 1996; Reznick et al. 1996; Zandona et al. 2011). In these prior analyses, maternal size was included as a covariate because guppies share an all but universal property of fish reproduction, which is that fecundity increases with body size. The goal was to ask if there are differences in fecundity between HP and LP guppies that are independent of maternal size. Here we replicated this analysis by including maternal weight as a covariate, which indeed showed that maternal-size-adjusted litter size was significantly smaller in LP guppies. However, we also analysed the data without the covariate. We did so to understand the effects of food levels on all life-history traits, and some of these effects are incorporated into maternal size because well-fed fish are larger at any given age. When size is not included as a covariate, food effects on litter size become more significant and habitat effects disappear.

### Traits unaffected by food levels and ancestral habitats

Thirteen traits (41%) were largely unaffected by food levels, including inter-birth intervals, offspring weights, the percentage fat in dry weights of both females and offspring, and the reproductive allotment. Interestingly, food-independent traits were only loosely embedded in the phenotypic correlation matrix. The absence of correlations eliminates potential indirect effects of food availability, further reducing the impact of resources on food-insensitive traits.

The large number of food-independent traits is somewhat surprising if traits associated with life histories are understood as complex, emergent properties of interactions between the entire phenotype and its (resource) environment (Coulson et al., 2006). It is worth noting that some of these traits are proportions (percentage fat, reproductive allotment). Food levels can have no effect on proportions in two broad ways: by truly affecting neither numerator nor denominator, or by affecting both of them proportionately so that the ratio itself stays constant. As some numerators (e.g., total dry weight of offspring in a female’s last litter) and denominators (e.g., total female dry weight) of ostensibly food-insensitive proportions were found to depend on food levels, our results do not necessarily contradict the notion that resources have pervasive effects on life histories.

Most food-independent traits also lacked significant habitat effects; only inter-birth intervals and offspring weights proved relatively unaffected by food levels yet differed among guppy ecotypes. Considering all the traits studied here, it appears that low resource availability impacts reproduction primarily by slowing down reproductive maturation rather than embryonic development, and by reducing the number of offspring rather than their size.

For the remaining traits, the putative reasons for the lack of food and habitat effects are diverse. For the percentage fat in females, we largely failed to see the previously documented (Reznick and Yang 1993) decrease in fat storage at low food. One reason for this may be that females compensated for low food levels by growing and maturing more slowly, but maintaining investment into fat reserves, as suggested by a study where low-food females had less body fat at maturity, but not anymore at parturition 3 (Auer 2010). Another reason may lie in the coarse nature of the ether extractions performed to remove stored fat. In addition to triglycerides, the main constituents of body fat, ether also dissolves other types of lipids, such as cholesterol, cholesterol esters, and free fatty acids (Cowey and Sargent 1972). If these other lipids were affected by the food treatment, or differed between ancestral habitats, effects on body fat might have been cancelled out. No effects of food or habitat were expected for the age and weight at the beginning of experiments: At that time, food manipulations had no yet begun, and habitat effects on weights, if present, might have been obscured by non-random variation in the age when fish were admitted to the experiment (range 19-40 days).

## Conclusions

Using the ecotypes of Trinidadian guppies as an example, we show that the traits underlying life-history strategies can substantially differ in the magnitude and direction of plastic and genetic components. Life histories may therefore not evolve as a single unit, but may be composed of traits with different sensitivities to environmental conditions and local selection pressures, and consequently with highly individual rates of phenotypic and evolutionary change. To broadly speak of “life-history evolution” thus likely masks the complex interplay of genes and environment on the multiple traits that underpin life-history strategies.

## Acknowledgements

This research was funded by a Swiss National Science Early Postdoc Mobility Fellowship (project no. P2EZP3_181775) to A.F.

## Author contributions

A.F., T.C. and D.N.R. conceived the study. D.N.R provided the datasets. A.F. analysed the data and drafted the manuscript, with input from all authors. All authors contributed to subsequent revisions.

## Data Availability Statement

The complete dataset will be available at the Dryad Digital Repository pending manuscript acceptance.

