## Supporting Information for "Life histories as mosaics: plastic and genetic components differ among traits that underpin life-history strategies"

#### Table of Contents

Supporting Methods ... 4
Supporting Methods 1: Datasets ... 4
Supporting Methods 2: Laboratory rearing protocol ... 4
Supporting Methods 3: Measurement of life-history traits ... 6
Supporting Methods 4: Differences in food levels between datasets ... 7
Supporting Methods 5: Effects of food level and ancestral habitat – datasets analysed jointly ... 9
Supporting Methods 6: Effects of food level and ancestral habitat – datasets analysed separately ... 11
Supporting Results: Differences in food levels between datasets ... 12
Supporting Figures ... 14
Figure S1. Map of sampling localities in northeastern Trinidad, West Indies. ... 14
Figure S2. Heat map of *p*-values of fixed effects, their interactions, and random effects in models fitted to phenotypic traits. ... 17
Figure S3. Drainage-specific effects of food levels and predation regime on female length at parturition 1 (len1). ... 18
Figure S4. Drainage-specific effects of food levels and predation regime on female length at parturition 3 (len3). ... 19
Figure S5. Drainage-specific effects of food levels and predation regime on male length at sexual maturity (lenmat). ... 20
Figure S6. Drainage-specific effects of food levels and predation regime on the mean dry weight of new-born offspring in litter 2 (mnemb2). ... 21
Figure S7. Drainage-specific effects of food levels and predation regime on female age at parturition 1 (agepart1). ... 22
Figure S8. Drainage-specific effects of food levels and predation regime on male age at sexual maturity (agemat). ... 23
Figure S9. Drainage-specific effects of food levels and predation regime on the mean percentage fat of new-born offspring in litter 2 (mnembfat2). ... 24
Figure S10. Network plots of phenotypic correlations between traits – when only including fish from HP localities and kept at high food levels. ... 25
Figure S11. Network plots of phenotypic correlations between traits – when only including fish from HP localities and kept at low food levels. ... 27

### Supporting Methods

#### Supporting Methods 1: Datasets

We here use four datasets of guppies that were subjected to a low- and a high-quantity food regime in the laboratory. Three datasets have previously been published fully or in part, while one is published here for the first time. All datasets include fish from multiple sampling localities, with all localities bar one being unique to a dataset. In combination, the datasets include 708 females and 470 males, derived from 16 localities situated in five drainages on the north slope and two drainages on the south slope of the Northern Range Mountains of Trinidad (see map in Fig. S1). The localities, years of collection of wild-caught females, and sample sizes are listed in Table S1. Dataset 1 uses fish from an introduction experiment where, in 1981, guppies from a HP locality ("Control") were transplanted to a LP locality ("Introduction") situated within the same drainage, and both localities were sampled four years post-introduction (for details see Reznick and Bryga, 1987). Dataset 2 includes paired HP and LP localities from three drainages. Four of these localities formed part of a larger survey of parallel evolution in the life histories of guppies inhabiting streams with different predation regimes (for details see Reznick and Bryga, 1996). Dataset 3 is unpublished. It contains three HP and three LP localities from four drainages; of these drainages, only one includes both a HP and a LP locality. Finally, dataset 4 consists of paired HP and LP localities situated in two drainages, and was created to study female senescence (for details see Reznick et al., 2004, Reznick et al., 2006). While datasets 1 to 3 contain both sexes, dataset 4 is female-only.

#### Supporting Methods 2: Laboratory rearing protocol

The laboratory methods used have been described in detail elsewhere (see, e.g., Reznick and Bryga, 1987). Briefly, in all datasets, experimental individuals consisted of the second generation of laboratory-born offspring derived from wild-caught, gravid females. Experimental individuals were kept in maternal sib-groups. Typically, all of the offspring from a given mother were reared together in two groups of five in two-gallon tanks that were next to each other on the shelf. When offspring were 26.2 ± 3.6 days old (mean ± SD, min: 19.0, max: 40.0), they were measured, weighed and sexed. Subsequently, two males and two females in each maternal sib-group were selected for the quantified feeding treatment, and one individual of each sex randomly assigned to a high or low food level, respectively.

The exact food levels used are unknown, but differed between datasets. In dataset 1, the low and high food levels were “systematically lower” (Reznick and Bryga, 1987, p. 1374) than in a previous study, in which food availability had been set at a level promoting 50 to 85% of the growth rate observed with *ad libitum* feeding (Reznick, 1983). In dataset 2, the high and low food levels were chosen to sustain 65-70% and 45-50% of the maximum growth rate, respectively (Reznick and Bryga, 1996). These levels had been established in a previous series of experiments, which also showed that uncontrolled food sources (e.g., algae) contribute minimally to growth (Reznick, 1980). No information is available about the food levels used in the unpublished dataset 3. In dataset 4, the food levels were the same as in dataset 2. All fish were fed liver paste in the morning, and a paste made of living brine shrimp nauplii (*Artemia* sp.) in the afternoon, with quantities being controlled volumetrically, to the nearest microlitre, using Hamilton micropipettes. In both food treatments, quantities of food were increased biweekly to accommodate growth.

The four tanks of each maternal sib-group, or two tanks in the case of dataset 4, which does not contain males, were kept together throughout the experiment. They were distributed around the laboratory in a stratified randomised block array to avoid confounding the laboratory microenvironment with sampling locality and experimental food levels. However, as a consequence, maternal identity is confounded with other factors, such as variation across the laboratory in temperature.

#### Supporting Methods 3: Measurement of life-history traits

Although some of the traits listed in the following may not usually be considered life-history traits (e.g., percentage fat), we here use the term in a wider sense to include both life-history and life-history-associated traits. For males, the measured phenotypic traits were age, wet weight and standard length at sexual maturity. For females, we measured age, wet weight, and standard length at parturitions 1, 2 and 3, the first and second inter-birth intervals (i.e., the time between parturitions 1 and 2, and between parturitions 2 and 3, respectively), the dry weight of and percentage fat in a female's somatic and reproductive tissues, the percentage fat in her total tissues, the reproductive allotment, the number of offspring in litters 1, 2 and 3, and the mean dry weight of and mean percentage fat in new-born offspring in litters 1, 2 and 3. As female size contributes to fecundity, we also analysed litter sizes when fitting the postpartum maternal wet weight as a covariate (following Reznick and Bryga, 1987); these maternal-weight-adjusted litter sizes are included as separate traits. In addition, we included the male and female age and wet weight at the beginning of the experiment, when the controlled feeding regime had not yet started, as negative controls for the effect of food.

We consequently collected data on 36 dependent variables. Only dataset 2 includes measurements of all variables; the numbers of variables available from datasets 1, 3 and 4 are 27, 24, and 25, respectively. Dataset 1 lacks all the variables pertaining to a female's third litter. Consequently, dataset 1 also differs from the other datasets inasmuch as the dry weights and percentage fat of females were measured after their second (not third) parturition, and the reproductive allotment calculated based on the second (not third) litter. In dataset 3, all post-mortem measurements of both females and offspring are lacking. Dataset 4 lacks the post-mortem measurements of females, and all the variables measured in males.

Male maturity was characterised by the development of the intromittent organ, the gonopodium, which was initially checked weekly, then daily as males approached maturity, until the attainment of sexual maturity. Females were mated once a week until first parturition, and then again within 24 hours after they gave birth to their first, second and third litters. The offspring in litters one to three were counted and preserved within 12 hours of birth. Females were preserved immediately after bearing their third litter, except in dataset 1, where females were preserved after two litters. Post mortem, females were dissected, the gut and gut contents discarded, and the somatic and reproductive tissues oven-dried overnight at 60°C. The tissues were then weighed separately to the nearest 0.1 mg (dry weights). After a series of ether extractions to remove fat deposits, tissues were weighed again (fat-free dry weights). We calculated the percentage fat in a female's total, somatic and reproductive tissues as the proportion of the dry weight that was lost after the ether extractions, i.e., as the difference between the dry and fat-free dry weights, divided by the dry weights. In the same way, we measured the mean dry weights of and mean percentage fat in offspring in litters 1 to 3. A female's reproductive allotment was computed by dividing the total dry weight of offspring in her last litter by the sum of her total dry weight and the litter's total dry weight.

#### Supporting Methods 4: Differences in food levels between datasets

We used analysis of variance with repeated measures to test for differences in female growth rates as a function of the experimental food level, the dataset, the ancestral habitat, and the drainage. The primary goal was to compare the chosen food levels among datasets by using growth as a proxy for the size of food rations. We focused on females because dataset 4 did not include males, and because females were weighed four times and males only twice.

Two analyses were performed, each on a different subset of the data. In the first, which included all four datasets, the dependent variable was the female wet weight at the beginning of the experiment, at parturition 1 and at parturition 2. The second analysis additionally included the female wet weight at parturition 3. This analysis only included datasets 2 to 4, as in dataset 1 females were euthanised after producing two litters and hence no data exist on their weight at parturition 3.

We analysed female weights with a repeated measures analysis of variance, using function “aov” in R (R Core Team, 2020). The predictors were food level, age category, dataset, all their interactions including the three-way-interaction, ancestral habitat, the interaction between food level and ancestral habitat, drainage, and maternal identity nested within drainage. Our formula contained individual identity as the single error term, to account for repeated measurements of females. The error term specified two different error strata, with appropriate models fitted within each stratum. We tested the effects of age and all its interactions within individuals, while we tested all the other effects between individuals. We compared models using raw, base-*e* log-transformed and square-root-transformed wet weights. Although the models gave very similar results, we found that a log-transformation resulted in the best fit, as judged from diagnostic plots.

After fitting the two full models, we computed pairwise comparisons to find out where the differences between datasets lay. Due to the strong interactions between datasets, food levels and age categories, each pairwise comparison of datasets was made within a given food level and age category, while using the full model residuals as the error term. This resulted in 36 pairwise comparisons in the first analysis (4 datasets, 3 age categories, 2 food levels), and 24 in the second analysis (3 datasets, 4 age categories, 2 food levels). In analyses of variance, the order in which predictors are fitted is important. To further investigate effects of ancestral habitats, we therefore ran additional analyses where habitat was fitted as the first predictor.

#### Supporting Methods 5: Effects of food level and ancestral habitat – datasets analysed jointly

To find appropriate models, we fitted three models per trait: one each where trait values were raw, base-*e* log-transformed, and square-root-transformed. To avoid undefined values for values of zero of six traits (dry weight of a female's reproductive tissues, percentage fat in a female's total and somatic tissues, mean percentage fat in new-born offspring in litters 1 to 3) when performing log transformations, 0.01 was added to all values of these traits before log-transforming them. For the number of offspring in litters 1, 2 and 3, we additionally fitted a generalised linear mixed model with Poisson errors. For traits that are proportions (e.g. percentage fat, reproductive allotment), we additionally fitted a GLMM with binomial errors. We then selected the best-fitting model based on diagnostic plots and model convergence. Only models that converged were selected.

All models included the experimental food treatment (high vs. low), the ancestral habitat (high- vs. low-predation), and the dataset (1 to 4) as categorical fixed effects. The reference level was high food, under high predation, using dataset 2, as only dataset 2 contained all traits. No interactions between fixed effects were fitted in models shown in the main text, but models including all two-way interactions between food levels, ancestral habitats, and datasets are provided here in the Supporting Information. In models with interactions we used the same data transformation (un-, log-, or square-root-transformed) that proved best in models without interactions. For three traits (litter sizes 1 and 2, mean dry weight of embryos in litter 2), the resulting models were singular or non-converging, and so a different data transformation was used (square-root, untransformed, square root, respectively) that allowed the model to converge.

For some traits, our models included covariates. We fitted additional analyses of litter sizes where the postpartum maternal wet weight was used as a covariate, and show these maternal-weight-adjusted analyses alongside analyses of raw litter sizes. We also evaluated the male wet weight at the beginning of experiments as a potential covariate for male standard length, wet weight, and age at sexual maturity (following Reznick and Bryga, 1987). Including the covariate did not change the model results, and so weight-unadjusted analyses of these traits are shown. Before fitting models including a covariate, covariates were mean-centred by subtracting the covariate's average from each individual value of the covariate. This was necessary to obtain intercepts comparable to those of models not containing covariates. Comparable intercepts were required for computing mean-standardised effect sizes.

As random effects we included maternal identity nested within drainage, provided that drainage had five levels or more. For eight traits we had data from four drainages only, and so drainage was fitted as a categorical fixed effect, with maternal identity as the sole random effect. Fitting maternal identity within locality was precluded by the strong collinearity between locality and predation regime. For two traits (male and female age at the beginning of the experiment), maternal identity could not be included as a random effect, as fish from the same mother came from a single litter, and thus were of the same age. Unlike the other datasets, the so-far unpublished dataset 3 contains four localities that are not paired HP and LP sites from a single drainage. The resulting imbalance of the data could potentially affect our results. We therefore repeated our analyses after excluding the unpaired localities for a subset of the traits, and found that the results were not substantially different. Consequently, we kept using the full dataset.

Outlier screening revealed three traits (age at parturition 3, inter-birth intervals 1 and 2) that showed moderately influential data points (maximum Cook's distance: 0.20), and additional models were fitted after excluding these outliers. The results of the outlier-free and original models were very similar, so outliers were retained in our final models.

#### Supporting Methods 6: Effects of food level and ancestral habitat – datasets analysed separately

We used the same type of data transformation that resulted in the best fit when analysing datasets jointly. Models included the experimental food treatment and the ancestral habitat as fixed effects, and maternal identity as a random effect, except for the age at the beginning of experiments, where maternal identity could not be included as all fish from a given mother were born on the same day. For datasets 2 and 4, containing 3 and 2 drainages, respectively, the drainage was included as a third fixed effect. For dataset 1, containing a single drainage, drainage was not included as a predictor. Dataset 3 contained four drainages, yet we could not include drainage as a predictor because its effects were confounded with those of the predation regime, with only one of the four drainages being represented by both a HP and a LP locality.

#### Supporting Results: Differences in food levels between datasets

In all datasets, females gained weight more slowly under low-food conditions (main effect of food level: $F_1 = 301.7$, $p < 0.00001$; age x food interaction: $F_2 = 102.7$, $p < 0.00001$; Tables S2 and S3). However, datasets differed from one another in terms of the mean weight of females (largest difference between datasets computed from raw data: 42% lower weight in dataset 1 vs. 3; main effect of dataset: $F_3 = 188.1$, $p < 0.00001$), the magnitude to which growth was reduced at the lower food level (range: 16.1% in dataset 3 vs. 33.3% in dataset 4; food x dataset interaction: $F_3 = 9.13$, $p < 0.00001$), and the shape of growth curves themselves (age x dataset interaction: $F_6 = 137.5$, $p < 0.00001$). When including all four ages at which females were weighed in datasets 2-4, datasets also differed in the extent to which the food levels resulted in progressively larger weight differences as fish aged (three-way interaction between food, age, and dataset: $F_6 = 7.92$, $p < 0.00001$).

Pairwise comparisons of datasets offered more insight into the effects of variation in food levels (Tables S2 and S3). Dataset 1 had the highest food levels at the start of experiments, but these levels increased very slowly, resulting in fish receiving substantially less food in this compared to other datasets. Dataset 2 had the second-lowest food levels after the start of experiments, followed by dataset 4, which additionally had a particularly pronounced contrast between the high and low food levels as females grew older. In the unpublished dataset 3, food levels were highest by far.

These results were corrected for the effects of drainage ($F_5 = 34.6$, $p < 0.00001$), maternal identity nested within drainage ($F_{343} = 1.8$, $p < 0.00001$), and type of ancestral habitat ($F_1 = 12.7$, $p = 0.00043$, Tables S2 and S3). While the former two effects were highly significant, differences between habitat types were less significant. Females were only 5.9% heavier when originating from LP rather than HP habitats ($F_1 = 12.7$, $p = 0.00043$). These results were very similar when we included all four weights per female, thus excluding dataset 1 (5.9% difference; $F_1$ = 19.4, $p$ = 0.00002), and when fitting habitat as the first predictor in either model ($F_1 \leq 15.8$, $p \geq 0.00009$).

#### Supporting Figures

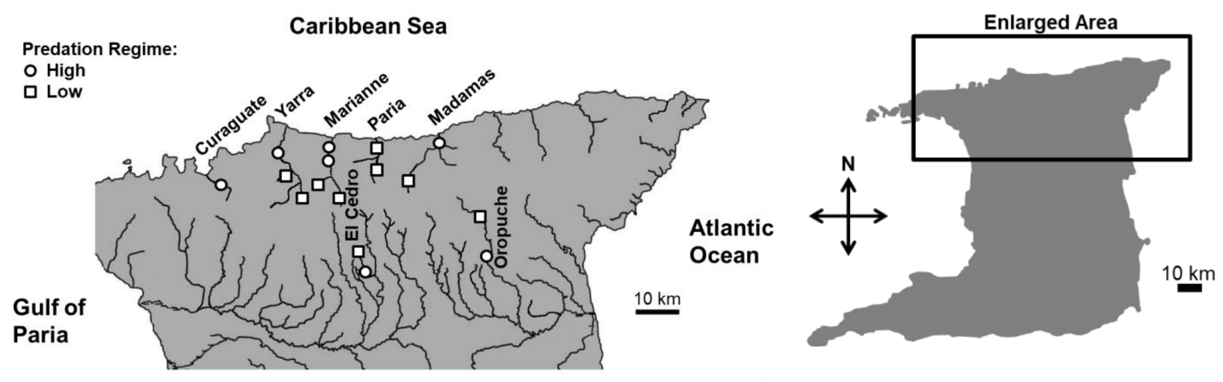

**Figure S1. Map of sampling localities in northeastern Trinidad, West Indies.**

Modified from Reznick and Travis (2019).

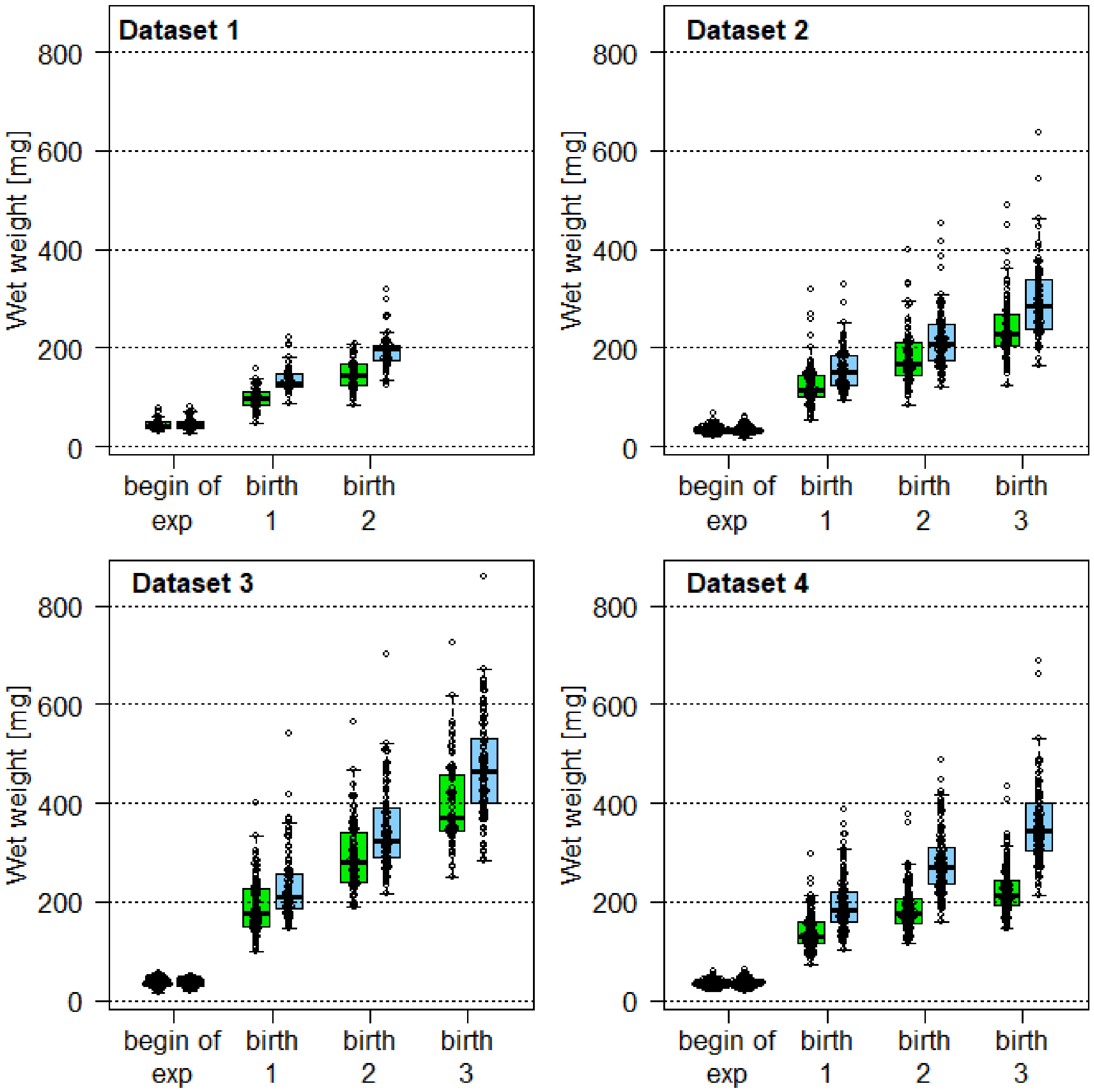

**Figure S2. Wet weight of female guppies in four datasets, illustrating consistent effects of experimental food treatments yet differences in food levels between datasets.**

Females were either subjected to a low-quantity diet (in green) or to a high-quantity diet (in blue), and were weighed at the beginning of the controlled food treatment (mean ± SD age: 26.8 ± 3.8 days), and when they gave birth for the first, second, and third time. Up to four data points may therefore stem from the same individual. For model results, see Tables S2 and S3.

| | Low food | Low predation | Dataset 1 | Dataset 3 | Dataset 4 | Food: Pred | Food: Data 1 | Food: Data 3 | Food: Data 4 | Pred: Data 1 | Pred: Data 3 | Pred: Data 4 | Maternal identity | Drainage | Covariate | |
|---|---|---|---|---|---|---|---|---|---|---|---|---|---|---|---|---|
| Group A | 0.00022 | 0.07 | 0.57 | 1.2e-18 | 0.04124 | 0.0239 | 0.02735 | 0.85 | 0.00829 | 0.53 | 0.26 | 0.25 | 0.05 | 9.12e-07 | | len1 |
| | 5.02e-07 | 0.23 | 0.61 | 1.64e-32 | 0.02347 | 0.17 | 0.0093 | 0.84 | 9.36e-08 | 0.27 | 0.02448 | 0.24 | 0.12 | 1.08e-09 | | len2 |
| | 6.07e-08 | 0.44 | | 1.48e-39 | 0.28 | 0.89 | | 0.96 | 2.66e-14 | | 5.62e-05 | 0.42 | 0.13 | 9.62e-12 | | len3 |
| | 2.74e-14 | 0.07 | | 2.59e-18 | | 0.88 | | 0.00013 | | | 0.41 | | 8.83e-10 | 0.08 | | lenmat |
| | 0.06 | 0.15 | 0.00983 | | | 0.07 | 0.04804 | | | 0.57 | | | 0.11 | 0.00833 * | | repwt |
| | 0.00012 | 0.09 | 0.00013 | | | 0.85 | 0.03171 | | | 0.02131 | | | 0.34 | 0.00239 * | | somwt |
| | 0.00434 | 0.07 | 0.3 | 0.00873 | 0.29 | 0.57 | 0.33 | 0.18 | 0.68 | 0.04971 | 0.66 | 0.6 | 1 | 0.49 | | n1 |
| | 0.00186 | 0.02727 | 0.87 | 3.6e-17 | 0.12 | 0.48 | 0.14 | 0.00191 | 0.69 | 0.01912 | 0.1 | 0.99 | 0.79 | 1 | | n2 |
| | 1.86e-07 | 0.03437 | | 3.23e-19 | 0.01791 | 0.74 | | 0.41 | 0.0138 | | 0.54 | 0.21 | 0.11 | 0.27 | | n3 |
| | 4.59e-09 | 0.00413 | 0.64 | 7.43e-21 | 0.00137 | 0.05 | 0.1 | 0.19 | 0.17 | 0.14 | 0.04746 | 0.25 | 0.01751 | 4.81e-14 | | wt1 |
| | 1.83e-07 | 0.00495 | 0.74 | 1.57e-38 | 5.25e-06 | 0.11 | 0.07 | 0.55 | 1.04e-06 | 0.07 | 0.0002 | 0.21 | 0.08 | 4.16e-16 | | wt2 |
| | 8.72e-11 | 0.07 | | 8.67e-44 | 0.00058 | 0.47 | | 0.34 | 8.32e-12 | | 2.63e-07 | 0.26 | 0.1 | 5.84e-19 | | wt3 |
| | 3.04e-26 | 0.18 | 0.7 | 1.16e-15 | | 0.04273 | 0.08 | 1.39e-07 | | 0.75 | 0.28 | | 2.71e-16 | 8.92e-06 | | wtmat |
| Group B | 0.59 | 0.03604 | 0.06 | 0.09 | 0.00062 | 0.13 | 0.91 | 0.58 | 0.19 | 0.03038 | 0.51 | 0.34 | 0.08 | 1 | | intrvl1 |
| | 0.00438 | 0.01743 | | 0.00849 | 0.00108 | 0.14 | | 0.00623 | 0.0215 | | 0.74 | 0.26 | 2.04e-15 | 0.01315 | | intrvl2 |
| | 0.62 | 0.00718 | 0.87 | | 0.00462 | 0.22 | 0.04007 | | 0.34 | 0.00445 | | 4.22e-07 | 0.01489 | 1.03e-18 | | mnemb1 |
| | 0.00296 | 0.00025 | 0.72 | | 0.15 | 0.89 | 0.99 | | 0.03064 | 0.00043 | | 3.65e-06 | 0.00662 | 1.63e-16 | | mnemb2 |
| | 0.11 | 0.07 | | | 0.27 | 0.09 | | | 0.02279 | | | 5.78e-05 | 1 | 2.39e-10* | | mnemb3 |
| | 0.01794 | 0.00074 | 0.64 | 0.01996 | 0.00757 | 0.42 | 0.43 | 0.82 | 0.01852 | 0.013 | 0.36 | 0.1 | 0.51 | 0.0009 | 3.78e-37 | n1_wt1adj |
| Group C | 0.00175 | 7.62e-05 | 0.84 | 0.15 | 0.00098 | 0.68 | 0.15 | 0.22 | 0.14 | 0.00225 | 0.00356 | 0.37 | 0.02044 | 0.00021 | 9.99e-30 | n2_wt2adj |
| | 6.76e-05 | 0.00365 | | 0.00051 | 0.00027 | 0.54 | | 0.66 | 0.65 | | 0.03408 | 0.09 | 0.05 | 6.03e-05 | 3.32e-18 | n3_wt3adj |
| | 2.74e-16 | 0.00011 | 0.58 | 0.81 | 0.48 | 0.00878 | 5.26e-08 | 7.62e-05 | 0.02433 | 0.24 | 0.5 | 0.38 | 0.00865 | 0.00069 | | agepart1 |
| | 2.7e-15 | 4.45e-05 | 0.99 | 0.56 | 0.1 | 0.06 | 7.39e-07 | 5.95e-05 | 0.04613 | 0.06 | 0.55 | 0.28 | 0.03756 | 0.00135 | | agepart2 |
| | 6.5e-14 | 0.00048 | | 0.29 | 0.02306 | 0.28 | | 1.75e-05 | 0.02405 | | 0.3 | 0.15 | 0.00107 | 0.00154 | | agepart3 |
| | 6.86e-07 | 0.05 | 0.38 | 0.01397 | | 0.00262 | 0.00813 | 0.00012 | | 0.0117 | 0.07 | | 8.2e-11 | 3.24e-08 | | agemat |
| Group D | 0.22 | 0.02537 | 6.62e-09 | | | 0.09 | 0.72 | | | 0.00837 | | | 0.89 | 0.31 * | | fat |
| | 0.79 | 0.37 | 0.00771 | | | 0.69 | 0.21 | | | 0.03386 | | | 1 | 0.06 * | | repfat |
| | 0.04861 | 0.04191 | 9.57e-09 | | | 0.25 | 0.91 | | | 0.01439 | | | 0.2 | 0.25 * | | somfat |
| | 0.4 | 0.6 | 0.49 | | 0.00696 | 0.18 | 0.88 | | 0.95 | 0.88 | | 0.63 | 1.17e-05 | 0.00013 | | mnembfat1 |
| | 0.24 | 0.63 | 0.61 | | 3.17e-05 | 0.68 | 0.54 | | 0.45 | 0.77 | | 0.52 | 3.18e-10 | 8.69e-07 | | mnembfat2 |
| | 0.07 | 0.78 | | | 1.06e-08 | 0.57 | | | 0.85 | | | 0.43 | 0.15 | 0.00104 * | | mnembfat3 |
| | 0.54 | 0.05 | 0.86 | | | 0.47 | 0.87 | | | 0.1 | | | 1 | 0.04902 * | | repall |
| | 0.97 | 0.46 | 0.04112 | 5.63e-09 | 2.25e-47 | 0.98 | 0.98 | 0.94 | 0.97 | 0.43 | 0.21 | 5.51e-06 | | 3.35e-14 | | age0 |
| | 0.86 | 0.47 | 0.00028 | 1.16e-11 | | 0.87 | 0.91 | 0.98 | | 0.52 | 0.04138 | | | 0.28 | | age0m |
| | 0.96 | 0.29 | 0.01019 | 0.01333 | 0.01441 | 0.34 | 0.24 | 0.92 | 0.01841 | 0.22 | 0.19 | 0.81 | 5.23e-69 | 4.98e-05 | | wt0 |
| | 0.25 | 0.68 | 0.05 | 5.62e-05 | | 0.01737 | 0.47 | 0.73 | | 0.46 | 0.07 | | 1.88e-60 | 0.00607 | | wt0m |

**Figure S3. Heat map of *p*-values of fixed effects, their interactions, and random effects in models fitted to phenotypic traits.**

A separate model was fitted for each of 36 traits. Models included the experimental food level (high vs. low), the ancestral habitat (high- vs. low-predation) and the dataset (1 to 4) as categorical fixed effects, with a reference level of high food, under high predation, using dataset 2. In addition, we fitted all the two-way interactions between fixed effects. Models of weight-adjusted litter size include postpartum maternal weight as a covariate. Random effects were maternal identity nested within drainage, unless there were only four drainages, in which case drainage was fitted as a categorical fixed effect. For these traits, the *p*-value of the overall effect of drainage comes from a log-likelihood ratio tests comparing the full model to one without drainage (indicated by an asterisk). In dataset 1 fish originated from a single drainage not sampled in any other dataset, so effects of dataset 1 and drainage are confounded. Traits were ordered as in Fig. 1. Effects were considered highly significant when $p < 0.000001$ (shaded in red), to account for an increased type I error rate due to multiple testing. Note that four traits in group D (age0, age0m, wt0, wt0m) were measured before controlled food treatments began and thus serve as negative controls for food effects. Trait abbreviations are as in Table 1.

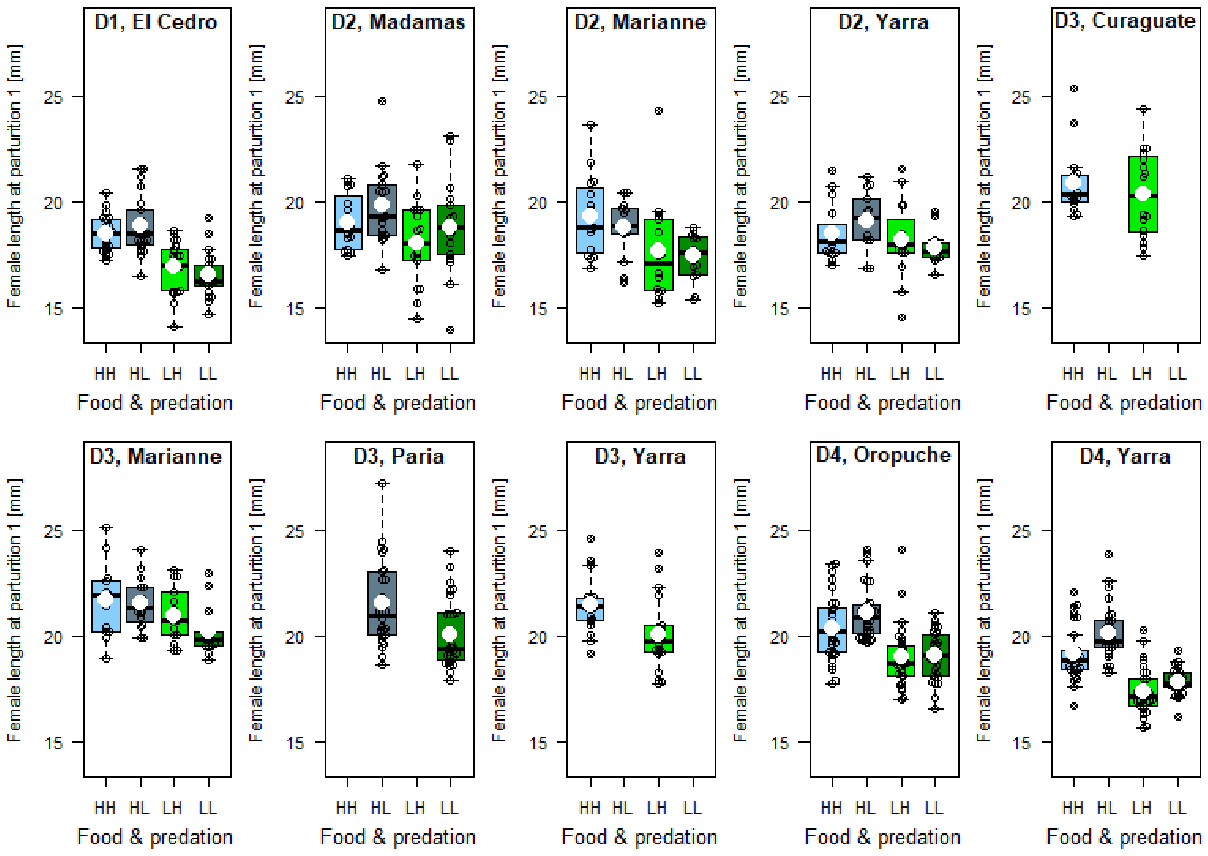

**Figure S4. Drainage-specific effects of food levels and predation regime on female length at parturition 1 (len1).**

This trait was assigned to group A (highly significant effect of food but not predation), but there is some variation to the overall pattern within drainages. White circles denote mean values of categorical predictor levels. D1-D4: dataset 1-4; HH: high food, high predation; HL: high food, low predation; LH: low food, high predation; LL: low food, low predation.

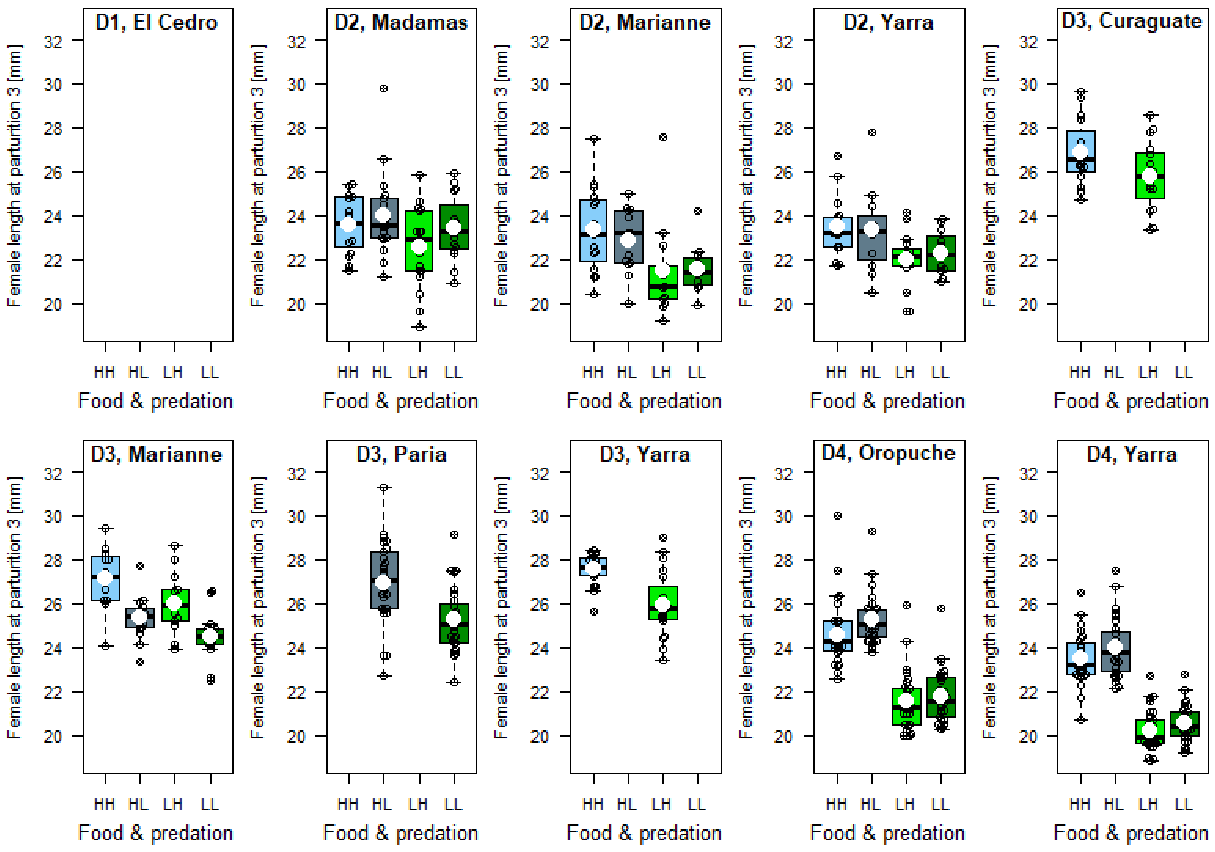

**Figure S5. Drainage-specific effects of food levels and predation regime on female length at parturition 3 (len3).**

This trait was assigned to group A (highly significant effect of food but not predation), but there is some variation to the overall pattern within drainages. White circles denote mean values of categorical predictor levels. The trait was not measured in dataset 1. D1-D4: dataset 1-4; HH: high food, high predation; HL: high food, low predation; LH: low food, high predation; LL: low food, low predation.

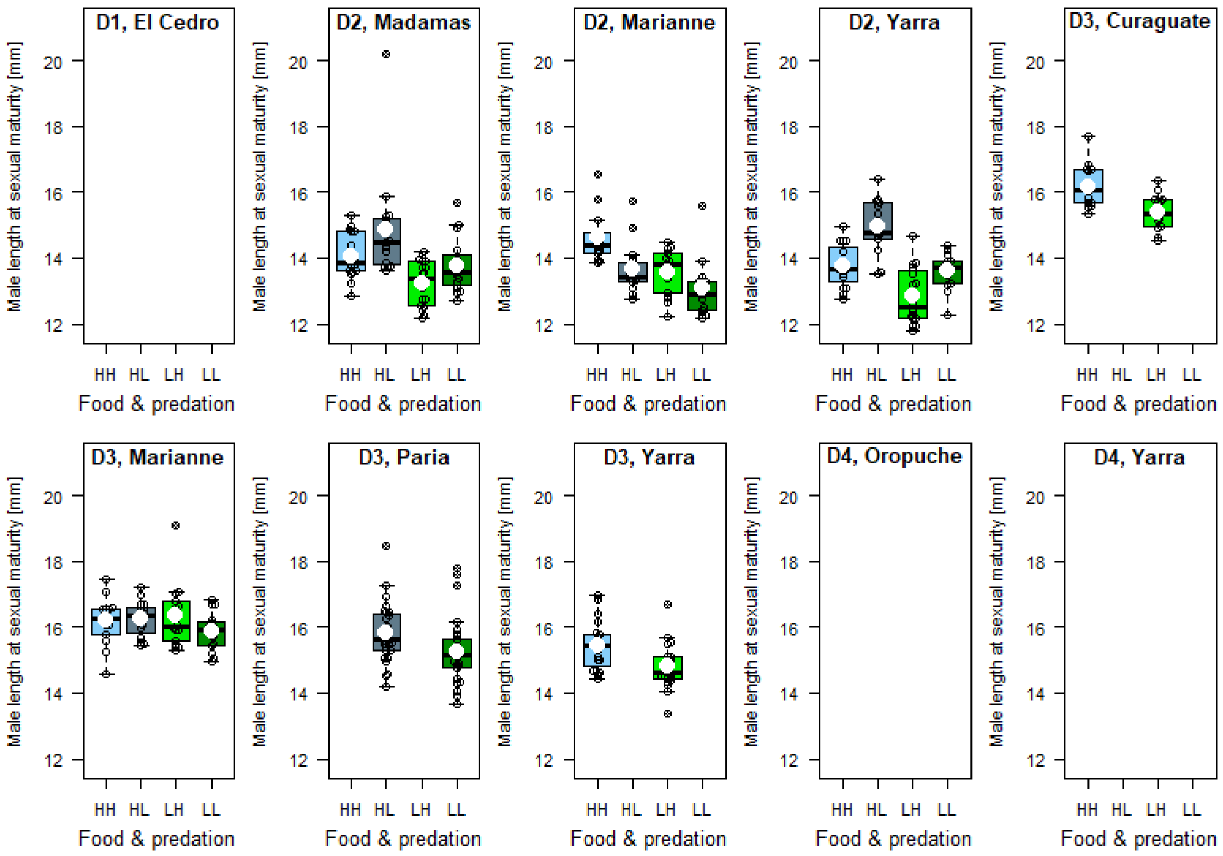

**Figure S6. Drainage-specific effects of food levels and predation regime on male length at sexual maturity (lenmat).**

This trait was assigned to group A (highly significant effect of food but not predation), but there is some variation to the overall pattern within drainages. White circles denote mean values of categorical predictor levels. The trait was not measured in datasets 1 and 4. D1-D4: dataset 1-4; HH: high food, high predation; HL: high food, low predation; LH: low food, high predation; LL: low food, low predation.

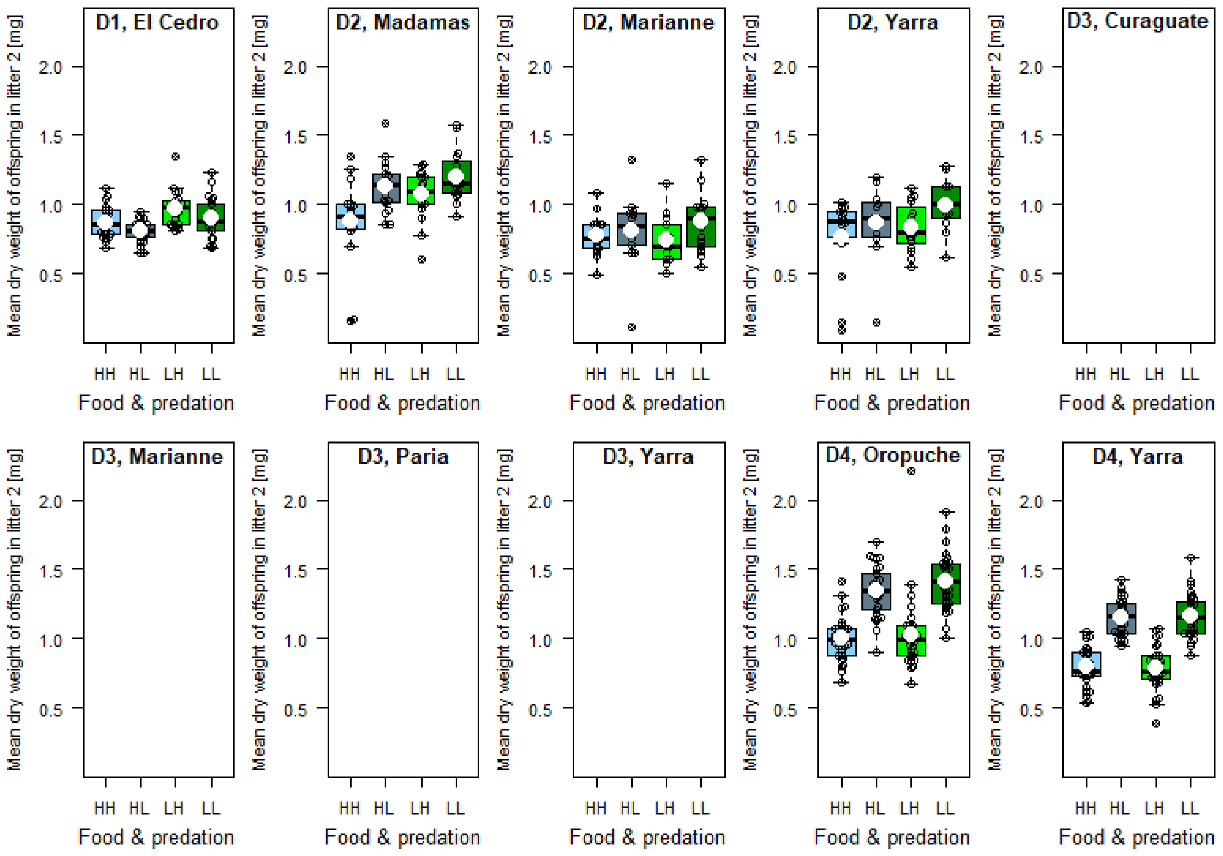

**Figure S7. Drainage-specific effects of food levels and predation regime on the mean dry weight of new-born offspring in litter 2 (mnemb2).**

This trait was assigned to group B (highly significant effect of predation but not food), but there is some variation to the overall pattern within drainages. White circles denote mean values of categorical predictor levels. The trait was not measured in dataset 3. D1-D4: dataset 1-4; HH: high food, high predation; HL: high food, low predation; LH: low food, high predation; LL: low food, low predation.

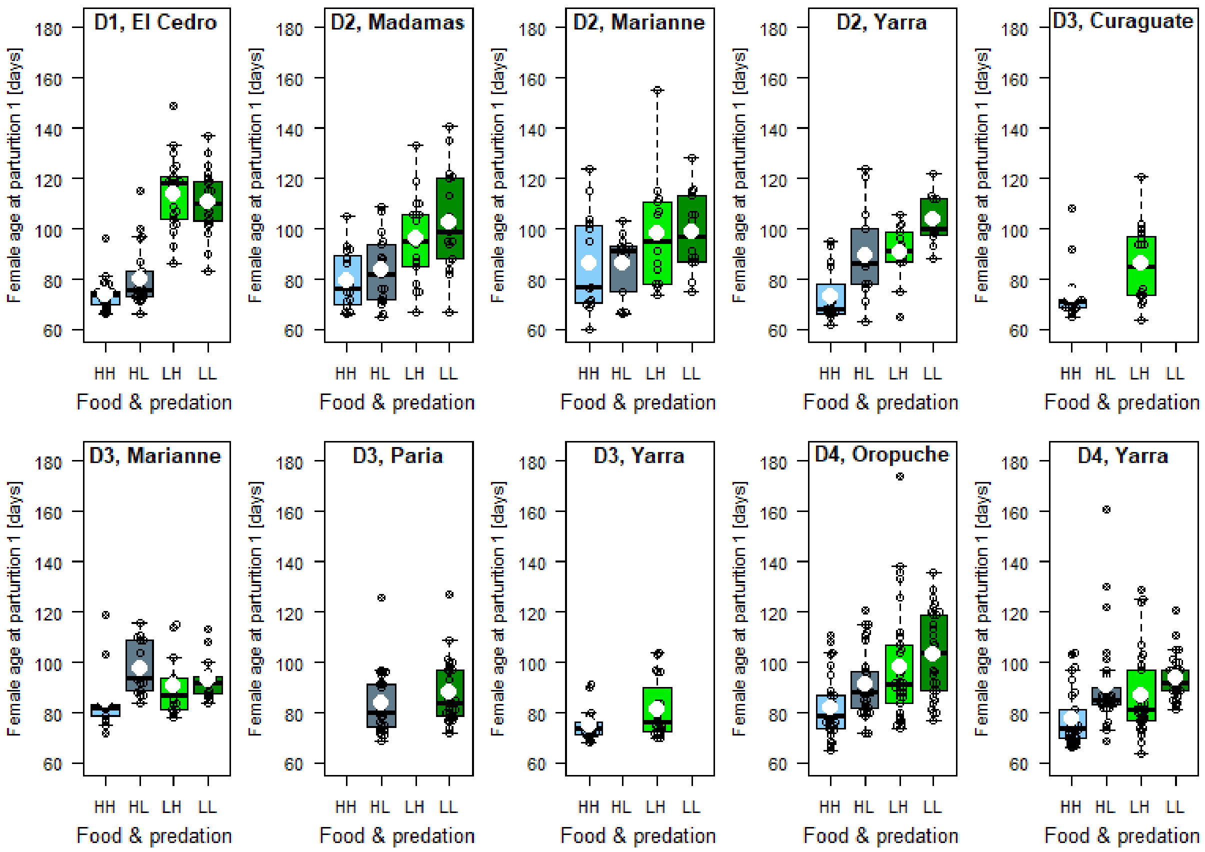

**Figure S8. Drainage-specific effects of food levels and predation regime on female age at parturition 1 (agepart1).**

This trait was assigned to group C (highly significant effect of both food and predation), but there is some variation to the overall pattern within drainages. White circles denote mean values of categorical predictor levels. D1-D4: dataset 1-4; HH: high food, high predation; HL: high food, low predation; LH: low food, high predation; LL: low food, low predation.

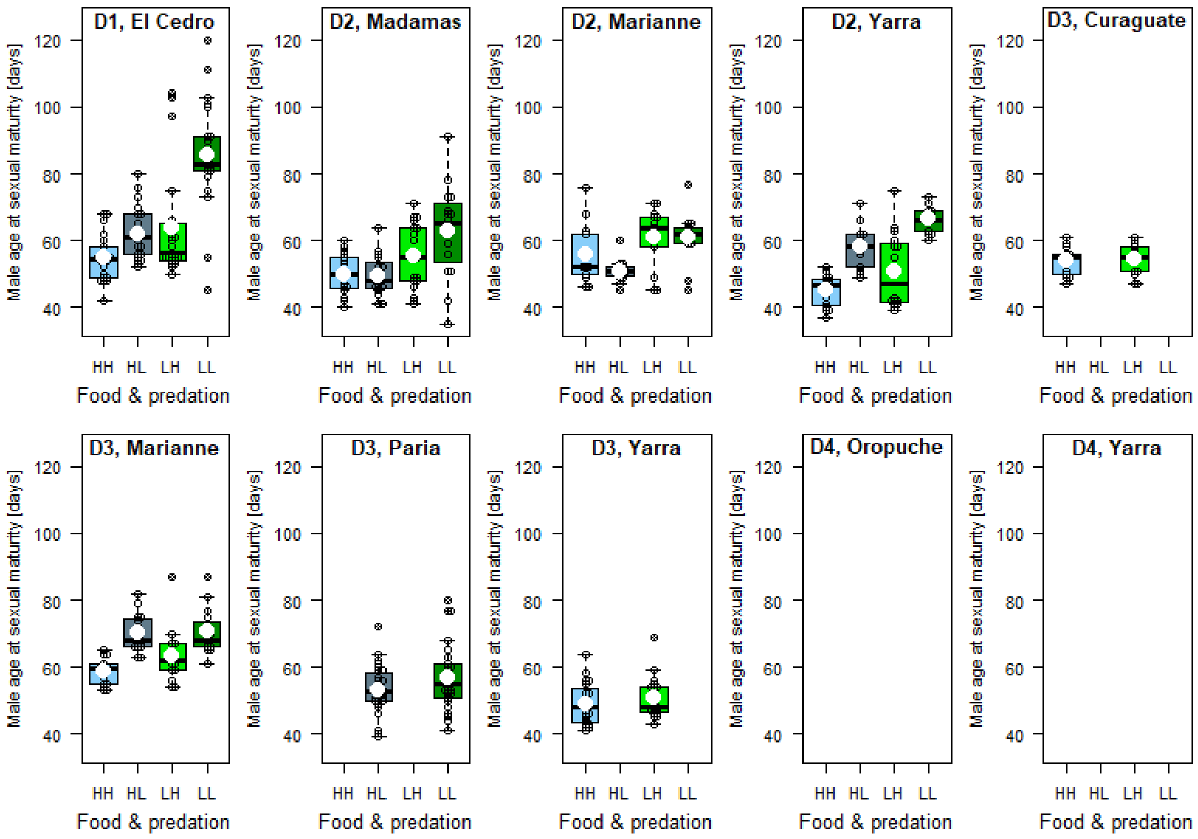

**Figure S9. Drainage-specific effects of food levels and predation regime on male age at sexual maturity (agemat).**

This trait was assigned to group C (highly significant effect of both food and predation), but there is some variation to the overall pattern within drainages. White circles denote mean values of categorical predictor levels. The trait was not measured in dataset 4. D1-D4: dataset 1-4; HH: high food, high predation; HL: high food, low predation; LH: low food, high predation; LL: low food, low predation.

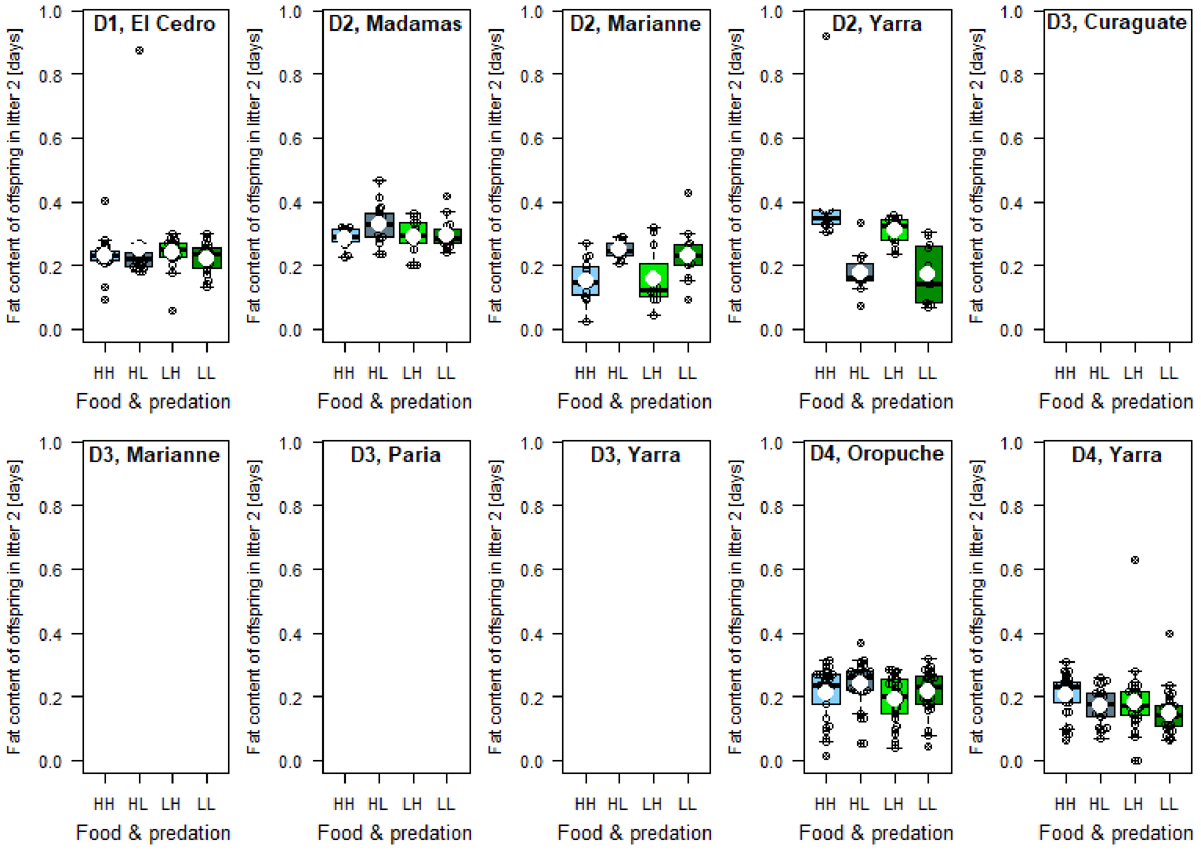

**Figure S10. Drainage-specific effects of food levels and predation regime on the mean percentage fat of new-born offspring in litter 2 (mnembfat2).**

This trait was assigned to group D (highly significant effect of neither food nor predation), but there is some variation to the overall pattern within drainages. White circles denote mean values of categorical predictor levels. The trait was not measured in dataset 3. D1-D4: dataset 1-4; HH: high food, high predation; HL: high food, low predation; LH: low food, high predation; LL: low food, low predation.

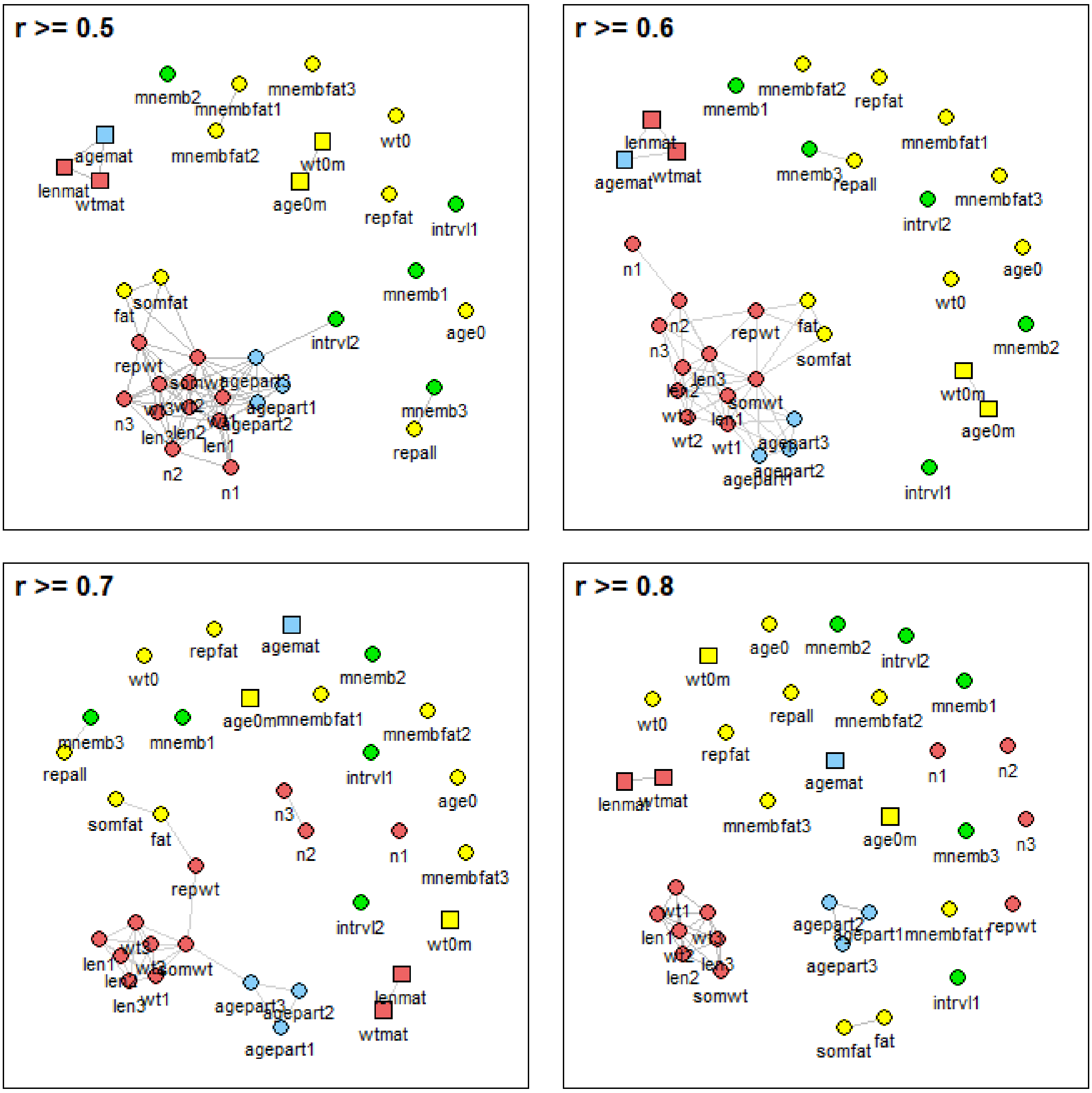

**Figure S11. Network plots of phenotypic correlations between traits – when only including fish from HP localities and kept at high food levels.**

Grey lines connect traits with pairwise Pearson product-moment correlation coefficients of $r \geq 0.5$, $\geq 0.6$, $\geq 0.7$ or $\geq 0.8$. No lines were plotted for correlation coefficients of $r < 0.5$. The correlation between traits measured in females (circles) and traits measured in males (squares) is $r = 0$ by definition. The position of traits and trait clusters relative to one another is irrelevant. Colours mark groups of traits with similar statistical dependence on experimental food levels and ancestral habitat, with group A (red) = highly significant effect of food but not

habitat, group B (green) = highly significant effect of habitat but not food, group C (blue) = highly significant effect of both, group D (yellow) = highly significant effect of neither. N = 300 individuals. Trait abbreviations are as in Table 1.

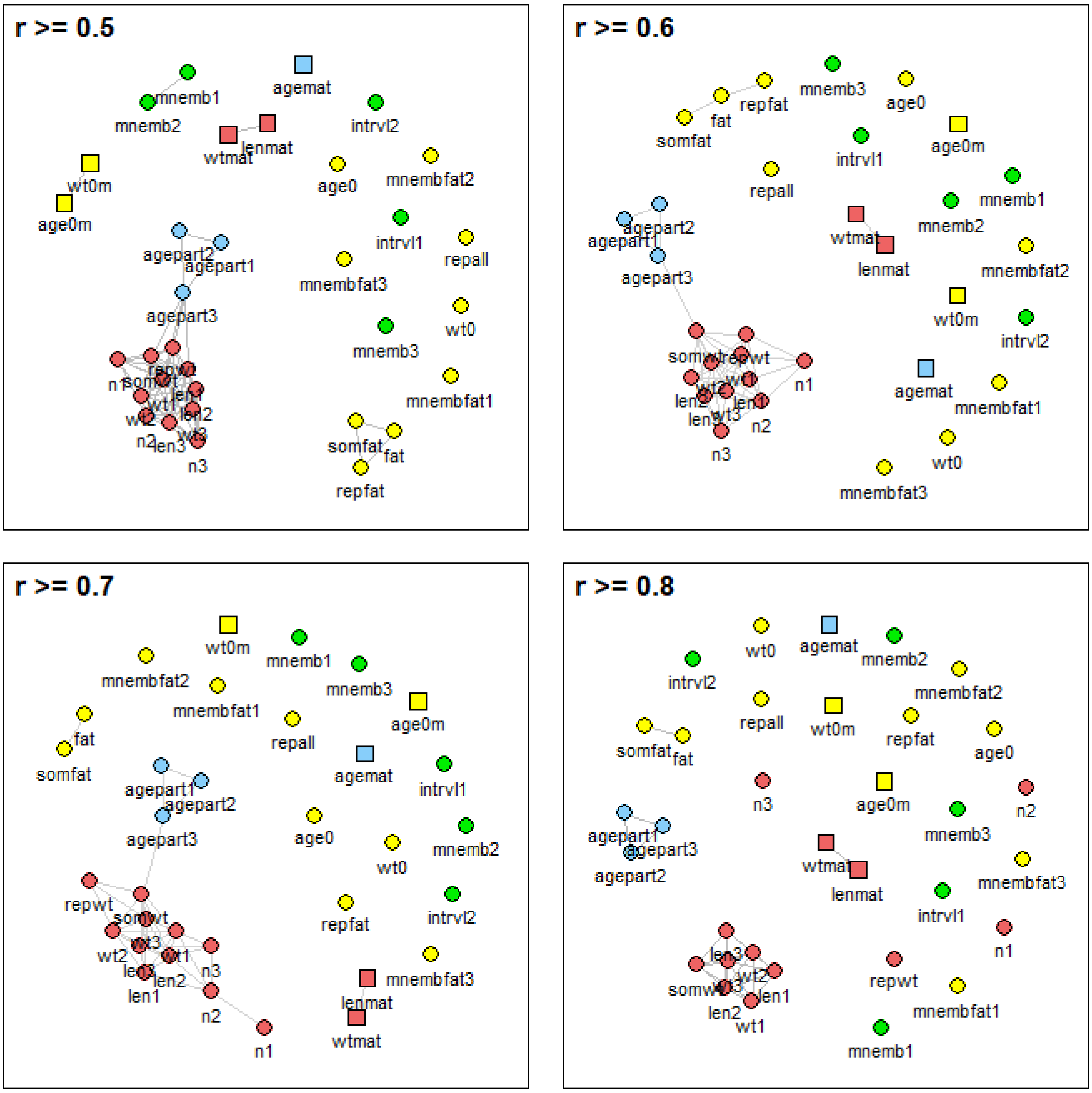

**Figure S12. Network plots of phenotypic correlations between traits – when only including fish from HP localities and kept at low food levels.**

Grey lines connect traits with pairwise Pearson product-moment correlation coefficients of $r \geq 0.5$, $\geq 0.6$, $\geq 0.7$ or $\geq 0.8$. No lines were plotted for correlation coefficients of $r < 0.5$. The correlation between traits measured in females (circles) and traits measured in males (squares) is $r = 0$ by definition. The position of traits and trait clusters relative to one another is irrelevant. Colours mark groups of traits with similar statistical dependence on experimental food levels and ancestral habitat, with group A (red) = highly significant effect of food but not

habitat, group B (green) = highly significant effect of habitat but not food, group C (blue) = highly significant effect of both, group D (yellow) = highly significant effect of neither. N = 302 individuals. Trait abbreviations are as in Table 1.

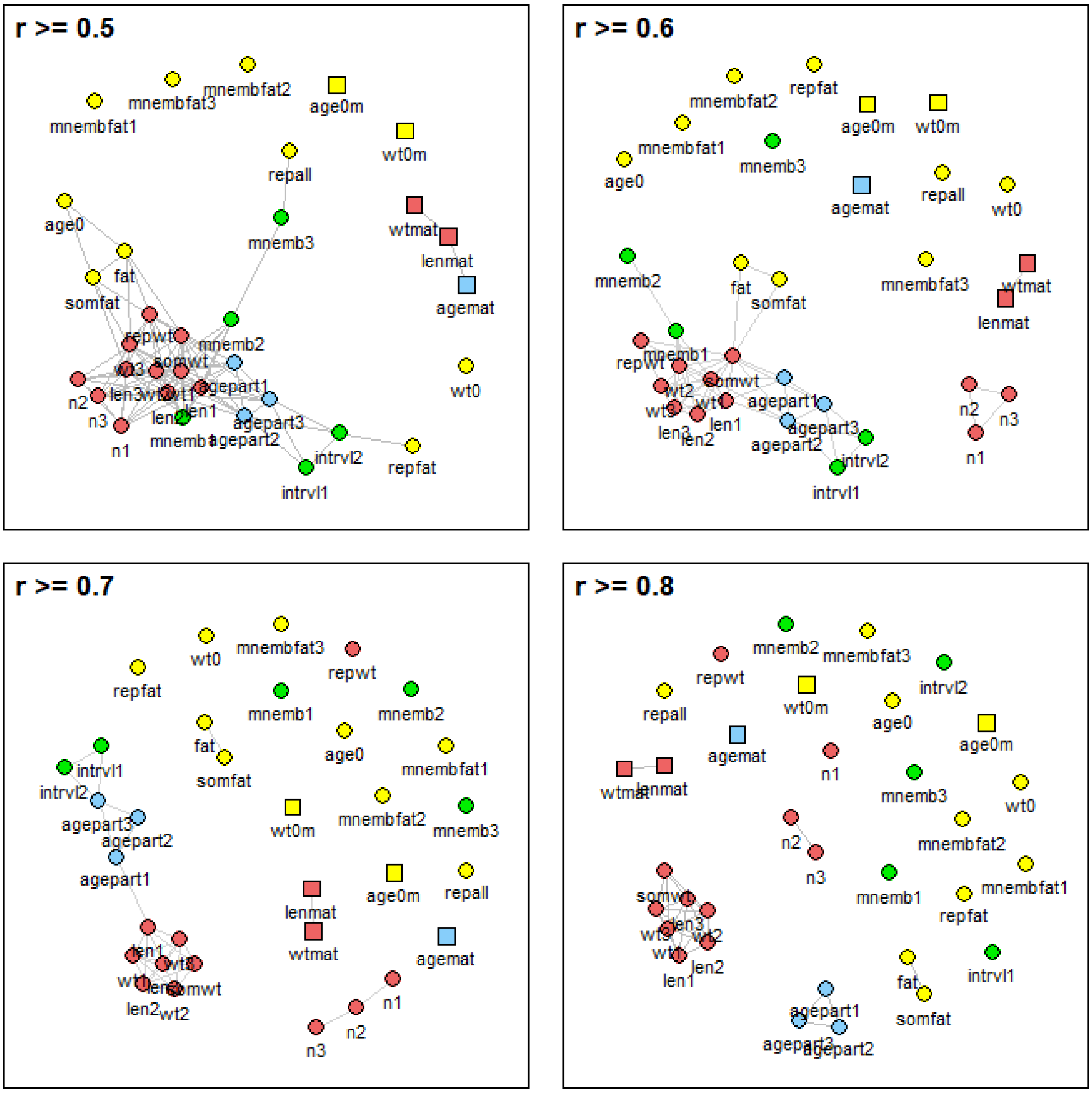

**Figure S13. Network plots of phenotypic correlations between traits – when only including fish from LP localities and kept at high food levels.**

Grey lines connect traits with pairwise Pearson product-moment correlation coefficients of $r \geq 0.5$, $\geq 0.6$, $\geq 0.7$ or $\geq 0.8$. No lines were plotted for correlation coefficients of $r < 0.5$. The correlation between traits measured in females (circles) and traits measured in males (squares) is $r = 0$ by definition. The position of traits and trait clusters relative to one another is irrelevant. Colours mark groups of traits with similar statistical dependence on experimental food levels and ancestral habitat, with group A (red) = highly significant effect of food but not

habitat, group B (green) = highly significant effect of habitat but not food, group C (blue) = highly significant effect of both, group D (yellow) = highly significant effect of neither. N = 287 individuals. Trait abbreviations are as in Table 1.

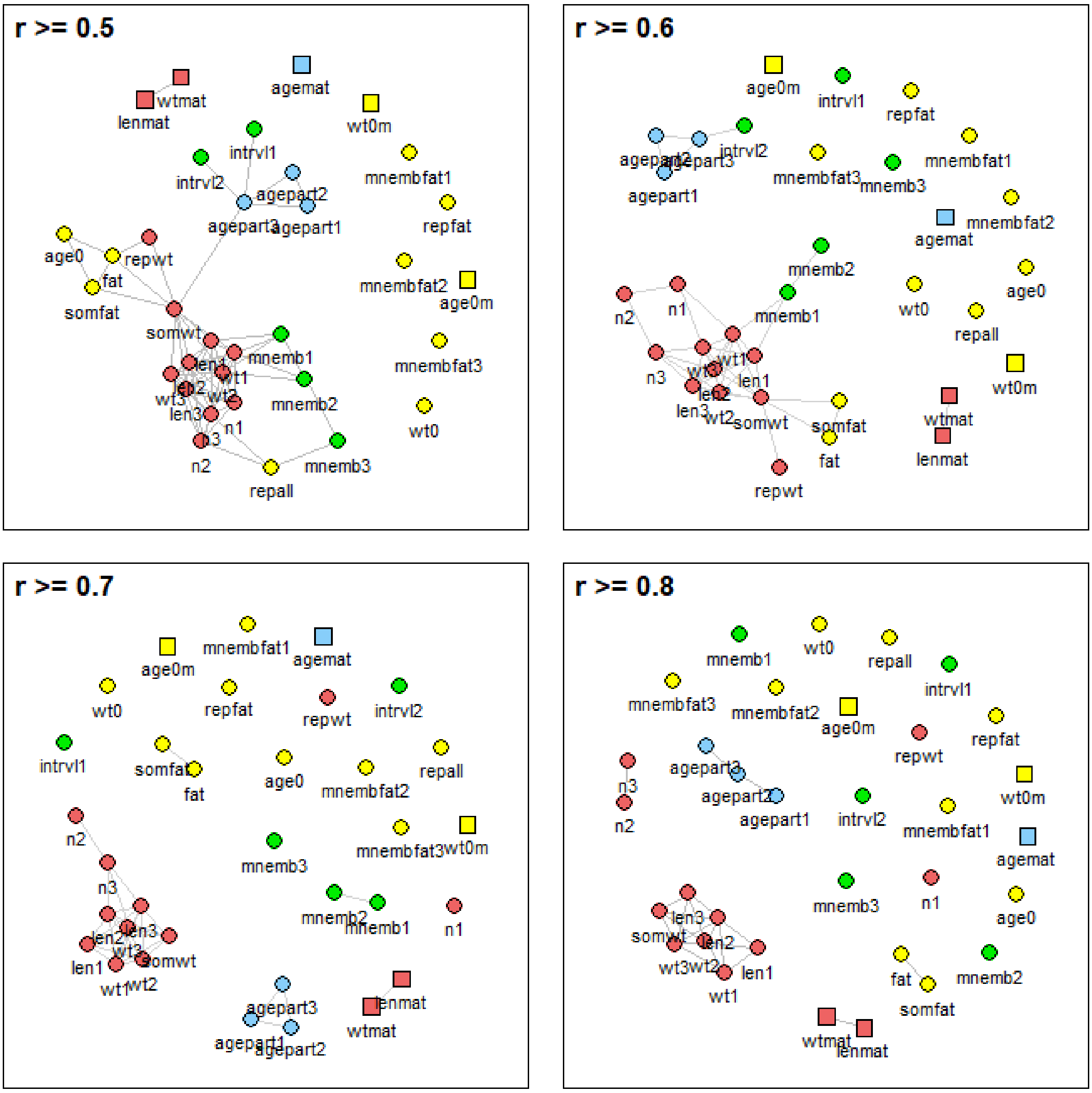

**Figure S14. Network plots of phenotypic correlations between traits – when only including fish from LP localities and kept at low food levels.**

Grey lines connect traits with pairwise Pearson product-moment correlation coefficients of $r \geq 0.5$, $\geq 0.6$, $\geq 0.7$ or $\geq 0.8$. No lines were plotted for correlation coefficients of $r < 0.5$. The correlation between traits measured in females (circles) and traits measured in males (squares) is $r = 0$ by definition. The position of traits and trait clusters relative to one another is irrelevant. Colours mark groups of traits with similar statistical dependence on experimental food levels and ancestral habitat, with group A (red) = highly significant effect of food but not

habitat, group B (green) = highly significant effect of habitat but not food, group C (blue) = highly significant effect of both, group D (yellow) = highly significant effect of neither. N = 288 individuals. Trait abbreviations are as in Table 1.

#### Supporting Tables

##### Table S1. Number of experimental fish per locality and dataset.

| Drainage | Locality | Habitat | Dataset | Year | Experimental fish | |
|---|---|---|---|---|---|---|
| | | | | | Females | Males |
| Curaguate (N) | Curaguate | HP | 3 | 1996 | 32 | 32 |
| El Cedro (S) | Control | HP | 1 | 1985 | 44 | 44 |
| | Introduction | LP | 1 | 1985 | 44 | 42 |
| Madamas (N) | Madamas | HP | 2 | 1991 | 36 | 36 |
| | Tapana-Aqui (tributary) | LP | 2 | 1991 | 34 | 34 |
| Marianne (N) | Marianne 1 | HP | 2 | 1990 | 30 | 30 |
| | Marianne 2 | HP | 3 | 1996 | 27 | 28 |
| | Marianne (tributary) | LP | 2 | 1990 | 31 | 32 |
| | Marianito (tributary) | LP | 3 | 1996 | 32 | 32 |
| Oropuche (S) | Oropuche | HP | 4 | 1998 | 60 | 0 |
| | Campo (tributary) | LP | 4 | 1998 | 60 | 0 |
| Paria (N) | Paria | LP | 3 | 1996 | 30 | 30 |
| | Paria (tributary) | LP | 3 | 1996 | 28 | 28 |
| Yarra (N) | Yarra | HP | 2 | 1990 | 32 | 32 |
| | | HP | 3 | 1996 | 40 | 40 |
| | | HP | 4 | 1998 | 60 | 0 |
| | Limon 1 (tributary) | LP | 2 | 1990 | 30 | 30 |
| | Limon 2 (tributary) | LP | 4 | 1998 | 58 | 0 |

The year of collection of field-caught ancestors is provided in brackets. Experimental fish are second-generation laboratory-reared offspring of wild-caught fish. Habitat: predation regime in ancestral stream; N and S: north and south slope of the Northern Range Mountains of Trinidad; HP: high predation; LP: low predation; f: number of female experimental fish; m: number of male experimental fish.

**Table S2. Analysis of variance with repeated measures of female growth until parturition 2 (wt0-wt2).**

| | Df | Sum Sq | Mean Sq | *F*-value | *p*-value |
|---|---|---|---|---|---|
| Error: between individuals | | | | | |
| Food | 1 | 17.64 | 17.64 | 301.66 | < 0.00001 |
| Dataset | 3 | 32.98 | 10.99 | 188.06 | < 0.00001 |
| Predation | 1 | 0.74 | 0.74 | 12.69 | 0.00043 |
| Drainage | 5 | 10.11 | 2.02 | 34.60 | < 0.00001 |
| Food x Dataset | 3 | 1.60 | 0.53 | 9.13 | < 0.00001 |
| Food x Predation | 1 | 0.15 | 0.15 | 2.60 | 0.11 |
| Drainage x Mother | 343 | 36.70 | 0.11 | 1.83 | < 0.00001 |
| Residuals | 315 | 18.41 | 0.06 | | |
| Error: within individuals | | | | | |
| Age | 2 | 1310.23 | 655.11 | 18348.95 | < 0.00001 |
| Age x Food | 2 | 7.33 | 3.67 | 102.71 | < 0.00001 |
| Age x Dataset | 6 | 29.45 | 4.91 | 137.46 | < 0.00001 |
| Age x Food x Dataset | 6 | 0.70 | 0.12 | 3.26 | 0.00345 |
| Residuals | 1330 | 47.49 | 0.04 | | |

Pairwise comparisons:

| | Mean ± SD Group 1 | Mean ± SD Group 2 | *F*-value | *p*-value |
|---|---|---|---|---|
| Error: within individuals | | | | |
| Food [h] Age [1] Data [1-2] | 45.3 ± 11.6 | 34.0 ± 7.9 | 91.20 | < 0.00001 |
| Food [h] Age [1] Data [1-3] | 45.3 ± 11.6 | 35.6 ± 7.3 | 60.09 | < 0.00001 |
| Food [h] Age [1] Data [1-4] | 45.3 ± 11.6 | 35.2 ± 7.6 | 75.43 | < 0.00001 |
| Food [h] Age [1] Data [2-3] | 34.0 ± 7.9 | 35.6 ± 7.3 | 4.06 | 0.04530 |
| Food [h] Age [1] Data [2-4] | 34.0 ± 7.9 | 35.2 ± 7.6 | 2.74 | 0.10 |
| Food [h] Age [1] Data [3-4] | 35.6 ± 7.3 | 35.2 ± 7.6 | 0.28 | 0.60 |
| Food [h] Age [2] Data [1-2] | 136.2 ± 24.7 | 156.9 ± 42.6 | 16.59 | < 0.00001 |
| Food [h] Age [2] Data [1-3] | 136.2 ± 24.7 | 231.7 ± 68.8 | 289.40 | < 0.00001 |
| Food [h] Age [2] Data [1-4] | 136.2 ± 24.7 | 195.0 ± 54.4 | 146.90 | < 0.00001 |

| | | | | |
|---|---|---|---|---|
| Food [h] Age [2] Data [2-3] | 156.9 ± 42.6 | 231.7 ± 68.8 | 244.90 | < 0.00001 |
| Food [h] Age [2] Data [2-4] | 156.9 ± 42.6 | 195.0 ± 54.4 | 90.69 | < 0.00001 |
| Food [h] Age [2] Data [3-4] | 231.7 ± 68.8 | 195.0 ± 54.4 | 59.35 | < 0.00001 |
| Food [h] Age [3] Data [1-2] | 196.0 ± 38.6 | 218.7 ± 59.5 | 10.01 | 0.00194 |
| Food [h] Age [3] Data [1-3] | 196.0 ± 38.6 | 348.0 ± 83.9 | 335.5 | < 0.00001 |
| Food [h] Age [3] Data [1-4] | 196.0 ± 38.6 | 278.6 ± 62.0 | 139.20 | < 0.00001 |
| Food [h] Age [3] Data [2-3] | 218.7 ± 59.5 | 348.0 ± 83.9 | 331.20 | < 0.00001 |
| Food [h] Age [3] Data [2-4] | 218.7 ± 59.5 | 278.6 ± 62.0 | 109.10 | < 0.00001 |
| Food [h] Age [3] Data [3-4] | 348.0 ± 83.9 | 278.6 ± 62.0 | 81.24 | < 0.00001 |
| Food [l] Age [1] Data [1-2] | 44.9 ± 10.9 | 34.0 ± 7.7 | 66.70 | < 0.00001 |
| Food [l] Age [1] Data [1-3] | 44.9 ± 10.9 | 35.7 ± 8.0 | 47.32 | < 0.00001 |
| Food [l] Age [1] Data [1-4] | 44.9 ± 10.9 | 33.7 ± 7.2 | 88.80 | < 0.00001 |
| Food [l] Age [1] Data [2-3] | 34.0 ± 7.7 | 35.7 ± 8.0 | 2.89 | 0.09 |
| Food [l] Age [1] Data [2-4] | 34.0 ± 7.7 | 33.7 ± 7.2 | 0.05 | 0.83 |
| Food [l] Age [1] Data [3-4] | 35.7 ± 8.0 | 33.7 ± 7.2 | 4.73 | 0.03090 |
| Food [l] Age [2] Data [1-2] | 97.6 ± 23.1 | 124.7 ± 43.5 | 43.29 | < 0.00001 |
| Food [l] Age [2] Data [1-3] | 97.6 ± 23.1 | 192.5 ± 56.1 | 430.30 | < 0.00001 |
| Food [l] Age [2] Data [1-4] | 97.6 ± 23.1 | 139.8 ± 37.6 | 148.6 | < 0.00001 |
| Food [l] Age [2] Data [2-3] | 124.7 ± 43.5 | 192.5 ± 56.1 | 277.9 | < 0.00001 |
| Food [l] Age [2] Data [2-4] | 124.7 ± 43.5 | 139.8 ± 37.6 | 30.89 | < 0.00001 |
| Food [l] Age [2] Data [3-4] | 192.5 ± 56.1 | 139.8 ± 37.6 | 185.00 | < 0.00001 |
| Food [l] Age [3] Data [1-2] | 145.9 ± 30.5 | 180.2 ± 54.2 | 27.08 | < 0.00001 |
| Food [l] Age [3] Data [1-3] | 145.9 ± 30.5 | 292.1 ± 68.1 | 389.40 | < 0.00001 |
| Food [l] Age [3] Data [1-4] | 145.9 ± 30.5 | 185.8 ± 44.2 | 56.64 | < 0.00001 |
| Food [l] Age [3] Data [2-3] | 180.2 ± 54.2 | 292.1 ± 68.1 | 275.90 | < 0.00001 |
| Food [l] Age [3] Data [2-4] | 180.2 ± 54.2 | 185.8 ± 44.2 | 2.89 | 0.09 |
| Food [l] Age [3] Data [3-4] | 292.1 ± 68.1 | 185.8 ± 44.2 | 312.5 | < 0.00001 |

The analysis includes female wet weights measured at the beginning of the experiment (wt0), at parturition 1 (wt1) and at parturition 2 (wt2), but not weights measured at parturition 3 (wt3) because those were lacking from one of the four datasets. We also excluded individuals with missing values. In total, the model included 2019 observations, 673 experimental individuals, 353 mothers of experimental individuals, and seven drainages. Wet weights were measured in milligrams and were base-*e* log-transformed before fitting the model. However, group-specific means and standard deviations for pairwise comparisons are provided as untransformed values for a better intuitive understanding of the direction and magnitude of differences between datasets. Df: degrees of freedom, Sum Sq: sum of squares, Mean Sq: mean squares, SD: standard deviation, h: high, l: low, Data: dataset.

**Table S3. Analysis of variance with repeated measures of female growth until parturition 3 (wt0-wt3).**

| | Df | Sum Sq | Mean Sq | *F*-value | *p*-value |
|---|---|---|---|---|---|
| Error: between individuals | | | | | |
| Food | 1 | 25.74 | 25.74 | 302.26 | < 0.00001 |
| Dataset | 2 | 48.77 | 24.39 | 286.40 | < 0.00001 |
| Predation | 1 | 1.65 | 1.65 | 19.35 | 0.00002 |
| Drainage | 5 | 12.64 | 2.53 | 29.69 | < 0.00001 |
| Food x Dataset | 2 | 3.71 | 1.85 | 21.77 | < 0.00001 |
| Food x Predation | 1 | 0.01 | 0.01 | 0.14 | 0.71 |
| Drainage x Mother | 298 | 42.37 | 0.14 | 1.67 | 0.00001 |
| Residuals | 264 | 22.48 | 0.09 | | |
| Error: within individuals | | | | | |
| Age | 3 | 1688.11 | 562.70 | 20519.21 | < 0.00001 |
| Age x Food | 3 | 7.17 | 2.39 | 87.10 | < 0.00001 |
| Age x Dataset | 6 | 14.40 | 2.40 | 87.51 | < 0.00001 |
| Age x Food x Dataset | 6 | 1.30 | 0.22 | 7.92 | < 0.00001 |
| Residuals | 1707 | 46.81 | 0.03 | | |

Pairwise comparisons:

| | Mean ± SD Group 1 | Mean ± SD Group 2 | *F*-value | p-value |
|---|---|---|---|---|
| Error: within individuals | | | | |
| Food [h] Age [1] Data [2-3] | 34.1 ± 7.9 | 35.6 ± 7.3 | 3.60 | 0.06 |
| Food [h] Age [1] Data [2-4] | 34.1 ± 7.9 | 35.3 ± 7.5 | 3.05 | 0.08 |
| Food [h] Age [1] Data [3-4] | 35.6 ± 7.3 | 35.3 ± 7.5 | 0.12 | 0.73 |
| Food [h] Age [2] Data [2-3] | 156.5 ± 42.7 | 231.4 ± 69.3 | 226.90 | < 0.00001 |
| Food [h] Age [2] Data [2-4] | 156.5 ± 42.7 | 196.1 ± 54.4 | 97.34 | < 0.00001 |
| Food [h] Age [2] Data [3-4] | 231.4 ± 69.3 | 196.1 ± 54.4 | 53.02 | < 0.00001 |
| Food [h] Age [3] Data [2-3] | 218.9 ± 59.8 | 346.8 ± 83.7 | 336.10 | < 0.00001 |
| Food [h] Age [3] Data [2-4] | 218.9 ± 59.8 | 280.2 ± 61.4 | 119.40 | < 0.00001 |
| Food [h] Age [3] Data [3-4] | 346.8 ± 83.7 | 280.2 ± 61.4 | 75.66 | < 0.00001 |

| | | | | |
|---|---|---|---|---|
| Food [h] Age [4] Data [2-3] | 294.7 ± 76.5 | 470.9 ± 102.5 | 301.30 | < 0.00001 |
| Food [h] Age [4] Data [2-4] | 294.7 ± 76.5 | 357.7 ± 77.6 | 64.94 | < 0.00001 |
| Food [h] Age [4] Data [3-4] | 470.9 ± 102.5 | 357.7 ± 77.6 | 118.30 | < 0.00001 |
| Food [l] Age [1] Data [2-3] | 34.0 ± 7.9 | 35.8 ± 8.1 | 3.15 | 0.08 |
| Food [l] Age [1] Data [2-4] | 34.0 ± 7.9 | 33.6 ± 7.2 | 0.13 | 0.72 |
| Food [l] Age [1] Data [3-4] | 35.8 ± 8.1 | 33.6 ± 7.2 | 3.97 | 0.01604 |
| Food [l] Age [2] Data [2-3] | 123.8 ± 43.6 | 190.4 ± 54.9 | 268.20 | < 0.00001 |
| Food [l] Age [2] Data [2-4] | 123.8 ± 43.6 | 140.4 ± 37.3 | 35.27 | < 0.00001 |
| Food [l] Age [2] Data [3-4] | 190.4 ± 54.9 | 140.4 ± 37.3 | 177.5 | < 0.00001 |
| Food [l] Age [3] Data [2-3] | 178.7 ± 53.9 | 290.6 ± 68.1 | 294.95 | < 0.00001 |
| Food [l] Age [3] Data [2-4] | 178.7 ± 53.9 | 186.3 ± 44.1 | 4.70 | 0.03140 |
| Food [l] Age [3] Data [3-4] | 290.6 ± 68.1 | 186.3 ± 44.1 | 314.80 | < 0.00001 |
| Food [l] Age [4] Data [2-3] | 242.6 ± 64.2 | 402.4 ± 84.3 | 281.50 | < 0.00001 |
| Food [l] Age [4] Data [2-4] | 242.6 ± 64.2 | 223.5 ± 49.7 | 6.91 | 0.00928 |
| Food [l] Age [4] Data [3-4] | 402.4 ± 84.3 | 223.5 ± 49.7 | 517.40 | < 0.00001 |

The analysis includes female wet weights measured at the beginning of the experiment (wt0), at parturition 1 (wt1), 2 (wt2), and 3 (wt3). As dataset 1 lacked weights measured at parturition 3, dataset 1 was excluded from this analysis. We also excluded individuals with missing values. In total, the model included 2300 observations, 575 experimental individuals, 307 mothers of experimental individuals, and six drainages. Wet weights were measured in milligrams and were base-*e* log-transformed before fitting the model. However, group-specific means and standard deviations for pairwise comparisons are provided as untransformed values for a better intuitive understanding of the direction and magnitude of differences between datasets. Df: degrees of freedom, Sum Sq: sum of squares, Mean Sq: mean squares, SD: standard deviation, h: high, l: low, Data: dataset.

**Table S4. Linear mixed-effects model on female age at the beginning of the experiment (age0).**

| Predictor | | | |
|---|---|---|---|
| Fixed effects: | Estimate (S.E.) | *t*-value | *p*-value |
| Intercept | 23.43 (0.67) | 35.16 | 4.04e-09 |
| Food (high vs. low) | -0.00 (0.22) | -0.01 | 1.00 |
| Predation (high vs. low) | -0.87 (0.24) | -3.68 | 0.00025 |
| Dataset (2 vs. 1) | 4.08 (1.60) | 2.55 | 0.05 |
| Dataset (2 vs. 3) | 2.77 (0.39) | 7.15 | 2.85e-12 |
| Dataset (2 vs. 4) | 7.69 (0.41) | 18.63 | 2.09e-60 |
| Random effects: | Var (S.D.) | $\chi^2_1$ | *p*-value |
| Drainage | 2.055 (1.434) | 55.54 | 9.17e-14 |
| Residual | 8.448 (2.907) | | |

The female age at the beginning of the experiment was measured in days and was fitted as untransformed values. The total number of observations included in the model is 707. Maternal identity could not be included as a random effect, as fish from the same mother came from a single litter, and thus were of the same age. S.E.: standard error, Var: variance, S.D.: standard deviation.

**Table S5. Linear mixed-effects model on male age at the beginning of the experiment (age0m).**

| Predictor | | | |
|---|---|---|---|
| Fixed effects: | Estimate (S.E.) | *t*-value | *p*-value |
| Intercept | 23.56 (0.38) | 61.78 | 2.70e-12 |
| Food (high vs. low) | -0.05 (0.24) | -0.23 | 0.82 |
| Predation (high vs. low) | -0.06 (0.26) | -0.22 | 0.83 |
| Dataset (2 vs. 1) | 3.53 (0.69) | 5.09 | 0.01194 |
| Dataset (2 vs. 3) | 2.92 (0.32) | 9.17 | 8.59e-15 |
| Random effects: | Var (S.D.) | $\chi^2_1$ | *p*-value |
| Drainage | 0.297 (0.545) | 2.51 | 0.11 |
| Residual | 6.460 (2.542) | | |

The male age at the beginning of the experiment was measured in days and was fitted as untransformed values. The total number of observations included in the model is 459. Maternal identity could not be included as a random effect, as fish from the same mother came from a single litter, and thus were of the same age. S.E.: standard error, Var: variance, S.D.: standard deviation.

**Table S6. Linear mixed-effects model on male age at sexual maturity (agemat).**

| Predictor | | | |
|---|---|---|---|
| Fixed effects: | Estimate (S.E.) | *t*-value | *p*-value |
| Intercept | 3.84 (0.04) | 95.67 | 4.32e-11 |
| Food (high vs. low) | 0.11 (0.01) | 9.40 | 6.98e-18 |
| Predation (high vs. low) | 0.14 (0.02) | 7.11 | 1.53e-11 |
| Dataset (2 vs. 1) | 0.20 (0.09) | 2.27 | 0.09 |
| Dataset (2 vs. 3) | 0.07 (0.02) | 2.76 | 0.00625 |
| Random effects: | Var (S.D.) | $\chi^2_1$ | *p*-value |
| Maternal identity | 0.009 (0.095) | 28.70 | 8.47e-08 |
| Drainage | 0.006 (0.075) | 28.31 | 1.03e-07 |
| Residual | 0.016 (0.127) | | |

The male age at sexual maturity was measured in days and was base-*e* log-transformed before fitting the model. Model results are provided on the transformed scale. The total number of observations included in the model is 442. S.E.: standard error, Var: variance, S.D.: standard deviation.

**Table S7. Linear mixed-effects model on female age at first birth (agepart1).**

| Predictor | | | |
|---|---|---|---|
| Fixed effects: | Estimate (S.E.) | *t*-value | *p*-value |
| Intercept | 4.37 (0.02) | 194.07 | 2.22e-24 |
| Food (high vs. low) | 0.15 (0.01) | 12.25 | 6.03e-29 |
| Predation (high vs. low) | 0.08 (0.01) | 6.31 | 9.14e-10 |
| Dataset (2 vs. 1) | 0.04 (0.04) | 0.94 | 0.39 |
| Dataset (2 vs. 3) | -0.04 (0.02) | -2.03 | 0.04387 |
| Dataset (2 vs. 4) | 0.01 (0.02) | 0.24 | 0.81 |
| Random effects: | Var (S.D.) | $\chi^2_1$ | *p*-value |
| Maternal identity | 0.001 (0.029) | 0.35 | 0.56 |
| Drainage | 0.001 (0.036) | 11.59 | 0.00066 |
| Residual | 0.024 (0.156) | | |

The female age at first birth was measured in days and was base-*e* log-transformed before fitting the model. Model results are provided on the transformed scale. The total number of observations included in the model is 687. S.E.: standard error, Var: variance, S.D.: standard deviation.

**Table S8. Linear mixed-effects model on female age at second birth (agepart2).**

| Predictor | | | |
|---|---|---|---|
| Fixed effects: | Estimate (S.E.) | *t*-value | *p*-value |
| Intercept | 4.62 (0.02) | 233.91 | 5.59e-22 |
| Food (high vs. low) | 0.13 (0.01) | 12.44 | 1.43e-29 |
| Predation (high vs. low) | 0.08 (0.01) | 7.18 | 5.12e-12 |
| Dataset (2 vs. 1) | 0.04 (0.04) | 1.11 | 0.32 |
| Dataset (2 vs. 3) | -0.03 (0.02) | -1.74 | 0.08 |
| Dataset (2 vs. 4) | 0.03 (0.02) | 1.82 | 0.07 |
| Random effects: | Var (S.D.) | $\chi^2_1$ | *p*-value |
| Maternal identity | 0.000 (0.022) | 0.22 | 0.64 |
| Drainage | 0.001 (0.033) | 9.72 | 0.00182 |
| Residual | 0.017 (0.132) | | |

The female age at second birth was measured in days and was base-*e* log-transformed before fitting the model. Model results are provided on the transformed scale. The total number of observations included in the model is 681. S.E.: standard error, Var: variance, S.D.: standard deviation.

**Table S9. Linear mixed-effects model on female age at third birth (agepart3).**

| Predictor | | | |
|---|---|---|---|
| Fixed effects: | Estimate (S.E.) | *t*-value | *p*-value |
| Intercept | 4.82 (0.02) | 244.22 | 5.29e-20 |
| Food (high vs. low) | 0.08 (0.01) | 9.36 | 2.11e-18 |
| Predation (high vs. low) | 0.09 (0.01) | 7.73 | 1.97e-13 |
| Dataset (2 vs. 3) | -0.01 (0.02) | -0.59 | 0.56 |
| Dataset (2 vs. 4) | 0.05 (0.02) | 2.80 | 0.00596 |
| Random effects: | Var (S.D.) | $\chi^2_1$ | *p*-value |
| Maternal identity | 0.003 (0.050) | 8.15 | 0.00430 |
| Drainage | 0.001 (0.034) | 9.54 | 0.00202 |
| Residual | 0.012 (0.109) | | |

The female age at third birth was measured in days and was base-*e* log-transformed before fitting the model. Model results are provided on the transformed scale. The total number of observations included in the model is 583. S.E.: standard error, Var: variance, S.D.: standard deviation.

**Table S10. Linear mixed-effects model on the percentage fat in a female's total tissues (fat).**

| Predictor | | | |
|---|---|---|---|
| Fixed effects: | Estimate (S.E.) | *t*-value | *p*-value |
| Intercept | 0.20 (0.01) | 31.52 | 9.18e-76 |
| Food (high vs. low) | -0.02 (0.00) | -3.83 | 0.00019 |
| Predation (high vs. low) | -0.00 (0.01) | -0.05 | 0.96 |
| Dataset (2 vs. 1) | -0.07 (0.01) | -10.67 | 1.21e-19 |
| Drainage (Madamas vs. Mar.) | 0.01 (0.01) | 1.40 | 0.16 |
| Drainage (Madamas vs. Yarra) | 0.00 (0.01) | 0.56 | 0.58 |
| Random effects: | Var (S.D.) | $\chi^2_1$ | *p*-value |
| Maternal identity | 0.000 (0.008) | 0.11 | 0.74 |
| Residual | 0.002 (0.040) | | |

The percentage fat in a female's total tissues is a proportion and fitted as untransformed values. The total number of observations included in the model is 260. As the analysis included data from only four drainages, drainage was fitted as a fixed effect. In dataset 1 fish originated from a single drainage (El Cedro), which was not sampled for any other dataset, so effects of dataset 1 and of the El Cedro drainage cannot be disentangled. S.E.: standard error, Mar: Marianne drainage, Var: variance, S.D.: standard deviation.

**Table S11. Linear mixed-effects model on the inter-birth interval 1 (intrvl1).**

| Predictor | | | |
|---|---|---|---|
| Fixed effects: | Estimate (S.E.) | *t*-value | *p*-value |
| Intercept | 3.09 (0.02) | 138.85 | 1.23e-16 |
| Food (high vs. low) | 0.04 (0.01) | 3.49 | 0.00054 |
| Predation (high vs. low) | 0.07 (0.01) | 5.18 | 4.33e-07 |
| Dataset (2 vs. 1) | 0.07 (0.04) | 1.61 | 0.20 |
| Dataset (2 vs. 3) | 0.03 (0.02) | 1.61 | 0.11 |
| Dataset (2 vs. 4) | 0.15 (0.02) | 6.42 | 7.81e-08 |
| Random effects: | Var (S.D.) | $\chi^2_1$ | *p*-value |
| Maternal identity | 0.003 (0.051) | 3.20 | 0.07 |
| Drainage | 0.001 (0.032) | 0.24 | 0.63 |
| Residual | 0.025 (0.157) | | |

The interval between a female's birth 1 and 2 was measured in days and was base-*e* log-transformed before fitting the model. Model results are provided on the transformed scale. The total number of observations included in the model is 680. S.E.: standard error, Var: variance, S.D.: standard deviation.

**Table S12. Linear mixed-effects model on the inter-birth interval 2 (intrvl2).**

| Predictor | | | |
|---|---|---|---|
| Fixed effects: | Estimate (S.E.) | *t*-value | *p*-value |
| Intercept | 3.08 (0.03) | 120.42 | 3.59e-16 |
| Food (high vs. low) | 0.02 (0.01) | 2.87 | 0.00442 |
| Predation (high vs. low) | 0.10 (0.02) | 6.31 | 1.20e-09 |
| Dataset (2 vs. 3) | 0.06 (0.02) | 2.55 | 0.01184 |
| Dataset (2 vs. 4) | 0.10 (0.02) | 4.04 | 0.00010 |
| Random effects: | Var (S.D.) | $\chi^2_1$ | *p*-value |
| Maternal identity | 0.010 (0.101) | 63.25 | 1.82e-15 |
| Drainage | 0.002 (0.044) | 6.90 | 0.00864 |
| Residual | 0.010 (0.099) | | |

The interval between a female's birth 2 and 3 was measured in days and was base-*e* log-transformed before fitting the model. Model results are provided on the transformed scale. The total number of observations included in the model is 582. S.E.: standard error, Var: variance, S.D.: standard deviation.

**Table S13. Linear mixed-effects model on female standard length at birth 1 (len1).**

| Predictor | | | |
|---|---|---|---|
| Fixed effects: | Estimate (S.E.) | *t*-value | *p*-value |
| Intercept | 19.20 (0.27) | 70.05 | 6.74e-13 |
| Food (high vs. low) | -1.48 (0.11) | -13.59 | 4.53e-34 |
| Predation (high vs. low) | 0.25 (0.13) | 1.97 | 0.04930 |
| Dataset (2 vs. 1) | -0.89 (0.61) | -1.46 | 0.21 |
| Dataset (2 vs. 3) | 2.56 (0.21) | 12.35 | 3.26e-27 |
| Dataset (2 vs. 4) | 0.37 (0.22) | 1.65 | 0.10 |
| Random effects: | Var (S.D.) | $\chi^2_1$ | *p*-value |
| Maternal identity | 0.209 (0.458) | 2.82 | 0.09 |
| Drainage | 0.277 (0.526) | 23.06 | 1.57e-06 |
| Residual | 2.029 (1.425) | | |

The female standard length at birth 1 was measured in millimetres and was fitted as untransformed values. The total number of observations included in the model is 687. S.E.: standard error, Var: variance, S.D.: standard deviation.

**Table S14. Linear mixed-effects model on female standard length at birth 2 (len2).**

| Predictor | | | |
|---|---|---|---|
| Fixed effects: | Estimate (S.E.) | *t*-value | *p*-value |
| Intercept | 3.07 (0.01) | 211.25 | 1.03e-14 |
| Food (high vs. low) | -0.09 (0.00) | -17.48 | 3.06e-49 |
| Predation (high vs. low) | 0.01 (0.01) | 0.92 | 0.36 |
| Dataset (2 vs. 1) | -0.05 (0.03) | -1.38 | 0.23 |
| Dataset (2 vs. 3) | 0.15 (0.01) | 16.54 | 4.59e-44 |
| Dataset (2 vs. 4) | 0.00 (0.01) | 0.00 | 1.00 |
| Random effects: | Var (S.D.) | $\chi^2_1$ | *p*-value |
| Maternal identity | 0.000 (0.014) | 0.55 | 0.46 |
| Drainage | 0.001 (0.030) | 35.28 | 2.85e-09 |
| Residual | 0.004 (0.065) | | |

The female standard length at birth 2 was measured in millimetres and was base-*e* log-transformed before fitting the model. Model results are provided on the transformed scale. The total number of observations included in the model is 678. S.E.: standard error, Var: variance, S.D.: standard deviation.

**Table S15. Linear mixed-effects model on female standard length at birth 3 (len3).**

| Predictor | | | |
|---|---|---|---|
| Fixed effects: | Estimate (S.E.) | *t*-value | *p*-value |
| Intercept | 4.89 (0.03) | 141.73 | 6.52e-14 |
| Food (high vs. low) | -0.22 (0.01) | -16.89 | 2.55e-45 |
| Predation (high vs. low) | 0.01 (0.01) | 0.42 | 0.67 |
| Dataset (2 vs. 3) | 0.37 (0.02) | 17.12 | 4.68e-45 |
| Dataset (2 vs. 4) | -0.08 (0.02) | -3.63 | 0.00034 |
| Random effects: | Var (S.D.) | $\chi_1^2$ | *p*-value |
| Maternal identity | 0.000 (0.012) | 0.00 | 0.96 |
| Drainage | 0.005 (0.072) | 40.58 | 1.89e-10 |
| Residual | 0.024 (0.154) | | |

The female standard length at birth 3 was measured in millimetres and was square-root-transformed before fitting the model. Model results are provided on the transformed scale. The total number of observations included in the model is 583. S.E.: standard error, Var: variance, S.D.: standard deviation.

**Table S16. Linear mixed-effects model on male standard length at sexual maturity (lenmat).**

| Predictor | | | |
|---|---|---|---|
| Fixed effects: | Estimate (S.E.) | *t*-value | *p*-value |
| Intercept | 2.64 (0.01) | 251.73 | 4.23e-18 |
| Food (high vs. low) | -0.05 (0.00) | -10.02 | 5.41e-19 |
| Predation (high vs. low) | 0.02 (0.01) | 1.87 | 0.06 |
| Dataset (2 vs. 3) | 0.13 (0.01) | 14.81 | 1.01e-23 |
| Random effects: | Var (S.D.) | $\chi^2_1$ | *p*-value |
| Maternal identity | 0.002 (0.039) | 31.54 | 1.96e-08 |
| Drainage | 0.000 (0.014) | 3.49 | 0.06 |
| Residual | 0.002 (0.046) | | |

The male standard length at sexual maturity was measured in millimetres and was base-*e* log-transformed before fitting the model. Model results are provided on the transformed scale. The total number of observations included in the model is 357. S.E.: standard error, Var: variance, S.D.: standard deviation.

**Table S17. Linear mixed-effects model on the mean dry weight of new-born offspring in litter 1 (mnemb1).**

| Predictor | | | |
|---|---|---|---|
| Fixed effects: | Estimate (S.E.) | *t*-value | *p*-value |
| Intercept | 0.82 (0.08) | 10.34 | 0.00125 |
| Food (high vs. low) | 0.07 (0.02) | 4.19 | 3.81e-05 |
| Predation (high vs. low) | 0.19 (0.02) | 9.07 | 3.36e-17 |
| Dataset (2 vs. 1) | -0.06 (0.17) | -0.37 | 0.74 |
| Dataset (2 vs. 4) | 0.00 (0.04) | -0.02 | 0.99 |
| Random effects: | Var (S.D.) | $\chi^2_1$ | *p*-value |
| Maternal identity | 0.011 (0.103) | 14.94 | 0.00011 |
| Drainage | 0.023 (0.151) | 63.40 | 1.68e-15 |
| Residual | 0.032 (0.180) | | |

The mean dry weight of new-born offspring in litter 1 was measured in milligrams and was fitted as untransformed values. The total number of observations included in the model is 501. S.E.: standard error, Var: variance, S.D.: standard deviation.

**Table S18. Linear mixed-effects model on the mean dry weight of new-born offspring in litter 2 (mnemb2).**

| Predictor | | | |
|---|---|---|---|
| Fixed effects: | Estimate (S.E.) | *t*-value | *p*-value |
| Intercept | 0.82 (0.07) | 11.01 | 0.00084 |
| Food (high vs. low) | 0.06 (0.02) | 3.72 | 0.00025 |
| Predation (high vs. low) | 0.21 (0.02) | 9.73 | 3.84e-19 |
| Dataset (2 vs. 1) | -0.06 (0.16) | -0.40 | 0.72 |
| Dataset (2 vs. 4) | 0.11 (0.04) | 3.00 | 0.00299 |
| Random effects: | Var (S.D.) | $\chi^2_1$ | *p*-value |
| Maternal identity | 0.015 (0.123) | 29.96 | 4.40e-08 |
| Drainage | 0.020 (0.141) | 61.55 | 4.31e-15 |
| Residual | 0.028 (0.166) | | |

The mean dry weight of new-born offspring in litter 2 was measured in milligrams and was fitted as untransformed values. The total number of observations included in the model is 494. S.E.: standard error, Var: variance, S.D.: standard deviation.

**Table S19. Linear mixed-effects model on the mean dry weight of new-born offspring in litter 3 (mnemb3).**

| Predictor | | | |
|---|---|---|---|
| Fixed effects: | Estimate (S.E.) | *t*-value | *p*-value |
| Intercept | 0.82 (0.04) | 21.97 | 3.10e-60 |
| Food (high vs. low) | 0.05 (0.03) | 1.83 | 0.07 |
| Predation (high vs. low) | 0.25 (0.03) | 9.33 | 2.11e-17 |
| Dataset (2 vs. 4) | 0.10 (0.04) | 2.23 | 0.02704 |
| Drainage (Madamas vs. Mar.) | -0.12 (0.05) | -2.43 | 0.01592 |
| Drainage (Madamas vs. Oro.) | 0.13 (0.06) | 2.21 | 0.02838 |
| Drainage (Madamas vs. Yarra) | -0.09 (0.05) | -1.90 | 0.06 |
| Random effects: | Var (S.D.) | $\chi^2_1$ | *p*-value |
| Maternal identity | 0.002 (0.047) | 0.07 | 0.79 |
| Residual | 0.065 (0.255) | | |

The mean dry weight of new-born offspring in litter 3 was measured in milligrams and was fitted as untransformed values. The total number of observations included in the model is 398. As the analysis included data from only four drainages, drainage was fitted as a fixed effect. S.E.: standard error, Mar: Marianne drainage, Oro: Oropuche drainage, Var: variance, S.D.: standard deviation.

**Table S20. Linear mixed-effects model on the mean percentage fat in new-born offspring in litter 1 (mnembfat1).**

| Predictor | | | |
|---|---|---|---|
| Fixed effects: | Estimate (S.E.) | *t*-value | *p*-value |
| Intercept | 0.25 (0.03) | 9.80 | 0.00080 |
| Food (high vs. low) | 0.00 (0.01) | 0.46 | 0.64 |
| Predation (high vs. low) | -0.01 (0.01) | -1.40 | 0.16 |
| Dataset (2 vs. 1) | -0.04 (0.05) | -0.74 | 0.52 |
| Dataset (2 vs. 4) | -0.06 (0.02) | -3.15 | 0.00200 |
| Random effects: | Var (S.D.) | $\chi^2_1$ | *p*-value |
| Maternal identity | 0.003 (0.053) | 18.77 | 1.47e-05 |
| Drainage | 0.002 (0.045) | 14.90 | 0.00011 |
| Residual | 0.007 (0.086) | | |

The mean percentage fat in new-born offspring in litter 1 is a proportion and was fitted as untransformed values. The total number of observations included in the model is 497. S.E.: standard error, Var: variance, S.D.: standard deviation.

**Table S21. Linear mixed-effects model on the mean percentage fat in new-born offspring in litter 2 (mnembfat2).**

| Predictor | | | |
| --- | --- | --- | --- |
| Fixed effects: | Estimate (S.E.) | *t*-value | *p*-value |
| Intercept | 0.52 (0.03) | 18.14 | 0.00014 |
| Food (high vs. low) | -0.02 (0.01) | -3.25 | 0.00133 |
| Predation (high vs. low) | -0.00 (0.01) | -0.34 | 0.73 |
| Dataset (2 vs. 1) | -0.03 (0.06) | -0.42 | 0.71 |
| Dataset (2 vs. 4) | -0.09 (0.02) | -5.24 | 4.43e-07 |
| Random effects: | Var (S.D.) | $\chi^2_1$ | *p*-value |
| Maternal identity | 0.003 (0.059) | 38.90 | 4.46e-10 |
| Drainage | 0.003 (0.053) | 24.43 | 7.70e-07 |
| Residual | 0.005 (0.070) | | |

The mean percentage fat in new-born offspring in litter 2 is a proportion and was square-root transformed before fitting the model. The total number of observations included in the model is 484. S.E.: standard error, Var: variance, S.D.: standard deviation.

**Table S22. Linear mixed-effects model on the mean percentage fat in new-born offspring in litter 3 (mnembfat3).**

| Predictor | | | |
|---|---|---|---|
| Fixed effects: | Estimate (S.E.) | *t*-value | *p*-value |
| Intercept | 0.58 (0.02) | 37.41 | 5.20e-104 |
| Food (high vs. low) | -0.04 (0.01) | -4.17 | 4.68e-05 |
| Predation (high vs. low) | -0.02 (0.01) | -1.98 | 0.04955 |
| Dataset (2 vs. 4) | -0.14 (0.02) | -7.93 | 1.31e-13 |
| Drainage (Madamas vs. Mar.) | -0.05 (0.02) | -2.65 | 0.00878 |
| Drainage (Madamas vs. Oro.) | 0.03 (0.02) | 1.13 | 0.26 |
| Drainage (Madamas vs. Yarra) | -0.01 (0.02) | -0.65 | 0.52 |
| Random effects: | Var (S.D.) | $\chi^2_1$ | *p*-value |
| Maternal identity | 0.001 (0.035) | 2.01 | 0.16 |
| Residual | 0.008 (0.091) | | |

The mean percentage fat in new-born offspring in litter 3 is a proportion and was square-root-transformed before fitting the model. The total number of observations included in the model is 379. As the analysis included data from only four drainages, drainage was fitted as a fixed effect. S.E.: standard error, Mar: Marianne drainage, Oro: Oropuche drainage, Var: variance, S.D.: standard deviation.

**Table S23. Linear mixed-effects model on the number of offspring in litter 1 (n1).**

| Predictor | | | |
|---|---|---|---|
| Fixed effects: | Estimate (S.E.) | *t*-value | *p*-value |
| Intercept | 1.26 (0.09) | 13.99 | 5.91e-07 |
| Food (high vs. low) | -0.39 (0.05) | -8.19 | 1.33e-15 |
| Predation (high vs. low) | -0.20 (0.05) | -4.04 | 6.22e-05 |
| Dataset (2 vs. 1) | -0.15 (0.18) | -0.84 | 0.46 |
| Dataset (2 vs. 3) | 0.27 (0.08) | 3.44 | 0.00079 |
| Dataset (2 vs. 4) | -0.11 (0.09) | -1.35 | 0.18 |
| Random effects: | Var (S.D.) | $\chi^2_1$ | *p*-value |
| Maternal identity | 0.000 (0.000) | 0.00 | 1.00 |
| Drainage | 0.022 (0.150) | 0.34 | 0.56 |
| Residual | 0.382 (0.618) | | |

The number of offspring in litter 1 was base-*e* log-transformed before fitting the model. Model results are provided on the transformed scale. The total number of observations included in the model is 687. S.E.: standard error, Var: variance, S.D.: standard deviation.

**Table S24. Linear mixed-effects model on the maternal-weight-adjusted number of offspring in litter 1 (n1_wt1adj).**

| Predictor | | | |
|---|---|---|---|
| Fixed effects: | Estimate (S.E.) | *t*-value | *p*-value |
| Intercept | 1.33 (0.11) | 12.08 | 7.85e-06 |
| Food (high vs. low) | -0.13 (0.05) | -2.85 | 0.00452 |
| Predation (high vs. low) | -0.28 (0.05) | -5.92 | 7.74e-09 |
| Dataset (2 vs. 1) | -0.02 (0.25) | -0.09 | 0.93 |
| Dataset (2 vs. 3) | -0.21 (0.08) | -2.51 | 0.01274 |
| Dataset (2 vs. 4) | -0.24 (0.08) | -2.91 | 0.00393 |
| Maternal weight | 0.01 (0.00) | 12.81 | 8.85e-34 |
| Random effects: | Var (S.D.) | $\chi^2_1$ | *p*-value |
| Maternal identity | 0.012 (0.110) | 0.45 | 0.50 |
| Drainage | 0.049 (0.222) | 8.75 | 0.00310 |
| Residual | 0.293 (0.542) | | |

The number of offspring in litter 1 was base-*e* log-transformed before fitting the model. Model results are provided on the transformed scale. To account for the contribution of female size to fecundity, we used the mean-centered postpartum maternal wet weight (wt1) as a covariate. The total number of observations included in the model is 684. S.E.: standard error, Var: variance, S.D.: standard deviation.

**Table S25. Linear mixed-effects model on the number of offspring in litter 2 (n2).**

| Predictor | | | |
|---|---|---|---|
| Fixed effects: | Estimate (S.E.) | *t*-value | *p*-value |
| Intercept | 2.78 (0.05) | 53.05 | 2.71e-203 |
| Food (high vs. low) | -0.51 (0.04) | -12.19 | 1.09e-28 |
| Predation (high vs. low) | -0.18 (0.04) | -4.07 | 5.95e-05 |
| Dataset (2 vs. 1) | 0.09 (0.08) | 1.18 | 0.24 |
| Dataset (2 vs. 3) | 0.91 (0.06) | 15.01 | 1.06e-39 |
| Dataset (2 vs. 4) | -0.21 (0.06) | -3.65 | 0.00030 |
| Random effects: | Var (S.D.) | $\chi^2_1$ | *p*-value |
| Maternal identity | 0.014 (0.118) | 0.50 | 0.48 |
| Drainage | 0.000 (0.000) | 0.00 | 1.00 |
| Residual | 0.301 (0.548) | | |

The number of offspring in litter 2 was square-root-transformed before fitting the model. Model results are provided on the transformed scale. The total number of observations included in the model is 682. S.E.: standard error, Var: variance, S.D.: standard deviation.

**Table S26. Linear mixed-effects model on the maternal-weight-adjusted number of offspring in litter 2 (n2_wt2adj).**

| Predictor | | | |
|---|---|---|---|
| Fixed effects: | Estimate (S.E.) | *t*-value | *p*-value |
| Intercept | 2.89 (0.09) | 31.80 | 1.92e-09 |
| Food (high vs. low) | -0.28 (0.04) | -6.36 | 4.97e-10 |
| Predation (high vs. low) | -0.25 (0.05) | -5.41 | 1.22e-07 |
| Dataset (2 vs. 1) | 0.16 (0.20) | 0.80 | 0.47 |
| Dataset (2 vs. 3) | 0.33 (0.09) | 3.80 | 0.00018 |
| Dataset (2 vs. 4) | -0.29 (0.08) | -3.64 | 0.00037 |
| Maternal weight | 0.00 (0.00) | 10.87 | 1.98e-25 |
| Random effects: | Var (S.D.) | $\chi^2_1$ | *p*-value |
| Maternal identity | 0.038 (0.195) | 6.05 | 0.01393 |
| Drainage | 0.028 (0.167) | 8.08 | 0.00447 |
| Residual | 0.231 (0.481) | | |

The number of offspring in litter 2 was square-root-transformed before fitting the model. Model results are provided on the transformed scale. To account for the contribution of female size to fecundity, we used the mean-centered postpartum maternal wet weight (wt2) as a covariate. The total number of observations included in the model is 675. S.E.: standard error, Var: variance, S.D.: standard deviation.

**Table S27. Linear mixed-effects model on the number of offspring in litter 3 (n3).**

| Predictor | | | |
|---|---|---|---|
| Fixed effects: | Estimate (S.E.) | *t*-value | *p*-value |
| Intercept | 3.62 (0.10) | 37.57 | 7.36e-11 |
| Food (high vs. low) | -0.67 (0.05) | -13.73 | 1.62e-33 |
| Predation (high vs. low) | -0.27 (0.06) | -4.77 | 3.03e-06 |
| Dataset (2 vs. 3) | 1.13 (0.08) | 13.47 | 5.25e-26 |
| Dataset (2 vs. 4) | -0.51 (0.09) | -5.67 | 1.50e-07 |
| Random effects: | Var (S.D.) | $\chi^2_1$ | *p*-value |
| Maternal identity | 0.032 (0.179) | 1.93 | 0.17 |
| Drainage | 0.026 (0.162) | 2.57 | 0.11 |
| Residual | 0.341 (0.584) | | |

The number of offspring in litter 3 was square-root-transformed before fitting the model. Model results are provided on the transformed scale. The total number of observations included in the model is 581. S.E.: standard error, Var: variance, S.D.: standard deviation.

**Table S28. Linear mixed-effects model on the maternal-weight-adjusted number of offspring in litter 3 (n3_wt3adj).**

| Predictor | | | |
|---|---|---|---|
| Fixed effects: | Estimate (S.E.) | *t*-value | *p*-value |
| Intercept | 3.68 (0.11) | 33.15 | 1.22e-09 |
| Food (high vs. low) | -0.41 (0.05) | -7.72 | 9.67e-14 |
| Predation (high vs. low) | -0.31 (0.06) | -5.53 | 7.19e-08 |
| Dataset (2 vs. 3) | 0.60 (0.10) | 6.05 | 4.49e-09 |
| Dataset (2 vs. 4) | -0.53 (0.09) | -5.91 | 1.56e-08 |
| Maternal weight | 0.00 (0.00) | 8.81 | 1.53e-17 |
| Random effects: | Var (S.D.) | $\chi^2_1$ | *p*-value |
| Maternal identity | 0.045 (0.212) | 4.72 | 0.02987 |
| Drainage | 0.045 (0.213) | 13.74 | 0.00021 |
| Residual | 0.285 (0.534) | | |

The number of offspring in litter 3 was square-root-transformed before fitting the model. Model results are provided on the transformed scale. To account for the contribution of female size to fecundity, we used the mean-centered postpartum maternal wet weight (wt3) as a covariate. The total number of observations included in the model is 581. S.E.: standard error, Var: variance, S.D.: standard deviation.

**Table S29. Linear mixed-effects model on the reproductive allotment (repall).**

| Predictor | | | |
|---|---|---|---|
| Fixed effects: | Estimate (S.E.) | *t*-value | *p*-value |
| Intercept | 0.11 (0.01) | 15.02 | 6.77e-37 |
| Food (high vs. low) | -0.00 (0.01) | -0.36 | 0.72 |
| Predation (high vs. low) | -0.01 (0.01) | -1.17 | 0.24 |
| Dataset (2 vs. 1) | 0.03 (0.01) | 3.65 | 0.00032 |
| Drainage (Madamas vs. Mar.) | 0.01 (0.01) | 1.32 | 0.19 |
| Drainage (Madamas vs. Yarra) | 0.02 (0.01) | 2.47 | 0.01401 |
| Random effects: | Var (S.D.) | $\chi_1^2$ | *p*-value |
| Maternal identity | 0.000 (0.000) | 0.00 | 1.00 |
| Residual | 0.002 (0.048) | | |

The reproductive allotment is a proportion and was fitted as untransformed values. The total number of observations included in the model is 259. As the analysis included data from only four drainages, drainage was fitted as a fixed effect. In dataset 1 fish originated from a single drainage (El Cedro), which was not sampled for any other dataset, so the effects of dataset 1 and of the El Cedro drainage cannot be disentangled. S.E.: standard error, Mar: Marianne drainage, Var: variance, S.D.: standard deviation.

**Table S30. Linear mixed-effects model on the percentage fat in a female's reproductive tissues (repfat).**

| Predictor | | | |
|---|---|---|---|
| Fixed effects: | Estimate (S.E.) | *t*-value | *p*-value |
| Intercept | -1.18 (0.02) | -47.69 | 1.25e-104 |
| Food (high vs. low) | 0.00 (0.02) | 0.10 | 0.92 |
| Predation (high vs. low) | -0.01 (0.02) | -0.48 | 0.63 |
| Dataset (2 vs. 1) | -0.10 (0.03) | -3.83 | 0.00020 |
| Drainage (Madamas vs. Mar.) | -0.04 (0.03) | -1.51 | 0.13 |
| Drainage (Madamas vs. Yarra) | 0.02 (0.03) | 0.81 | 0.42 |
| Random effects: | Var (S.D.) | $\chi^2_1$ | *p*-value |
| Maternal identity | 0.001 (0.025) | 0.01 | 0.91 |
| Residual | 0.025 (0.159) | | |

The percentage fat in a female's reproductive tissues is a proportion and was base-*e* log-transformed before fitting the model. Model results are provided on the transformed scale. The total number of observations included in the model is 255. As the analysis included data from only four drainages, drainage was fitted as a fixed effect. In dataset 1 fish originated from a single drainage (El Cedro), which was not sampled for any other dataset, so the effects of dataset 1 and of the El Cedro drainage cannot be disentangled. S.E.: standard error, Mar: Marianne drainage, Var: variance, S.D.: standard deviation.

**Table S31. Linear mixed-effects model on the dry weight of a female's reproductive tissues (repwt).**

| Predictor | | | |
|---|---|---|---|
| Fixed effects: | Estimate (S.E.) | *t*-value | *p*-value |
| Intercept | 3.48 (0.12) | 29.86 | 9.92e-71 |
| Food (high vs. low) | -0.52 (0.09) | -6.05 | 1.30e-08 |
| Predation (high vs. low) | 0.01 (0.10) | 0.15 | 0.88 |
| Dataset (2 vs. 1) | -0.91 (0.13) | -7.09 | 7.15e-11 |
| Drainage (Madamas vs. Mar) | 0.27 (0.14) | 1.91 | 0.06 |
| Drainage (Madamas vs. Yarra) | -0.18 (0.14) | -1.25 | 0.21 |
| Random effects: | Var (S.D.) | $\chi^2_1$ | *p*-value |
| Maternal identity | 0.072 (0.268) | 1.77 | 0.18 |
| Residual | 0.476 (0.690) | | |

The dry weight of a female's reproductive tissues was measured in milligrams and was square-root-transformed before fitting the model. Model results are provided on the transformed scale. The total number of observations included in the model is 262. As the analysis included data from only four drainages, drainage was fitted as a fixed effect. In dataset 1 fish originated from a single drainage (El Cedro), which was not sampled for any other dataset, so the effects of dataset 1 and the El Cedro drainage cannot be disentangled. S.E.: standard error, Mar: Marianne drainage, Var: variance, S.D.: standard deviation.

**Table S32. Linear mixed-effects model on the percentage fat in a female's somatic tissues (somfat).**

| Predictor | | | |
|---|---|---|---|
| Fixed effects: | Estimate (S.E.) | *t*-value | *p*-value |
| Intercept | 0.17 (0.01) | 25.99 | 4.37e-62 |
| Food (high vs. low) | -0.02 (0.00) | -4.19 | 4.98e-05 |
| Predation (high vs. low) | 0.00 (0.01) | 0.32 | 0.75 |
| Dataset (2 vs. 1) | -0.08 (0.01) | -10.37 | 7.05e-19 |
| Drainage (Madamas vs. Mar.) | 0.01 (0.01) | 1.48 | 0.14 |
| Drainage (Madamas vs. Yarra) | 0.00 (0.01) | 0.19 | 0.85 |
| Random effects: | Var (S.D.) | $\chi^2_1$ | *p*-value |
| Maternal identity | 0.000 (0.016) | 2.30 | 0.13 |
| Residual | 0.002 (0.039) | | |

The percentage fat in a female's somatic tissues is a proportion and was fitted as untransformed values. The total number of observations included in the model is 262. As the analysis included data from only four drainages, drainage was fitted as a fixed effect. In dataset 1 fish originated from a single drainage (El Cedro), which was not sampled for any other dataset, so the effects of dataset 1 and of the El Cedro drainage cannot be disentangled. S.E.: standard error, Mar: Marianne drainage, Var: variance, S.D.: standard deviation.

**Table S33. Linear mixed-effects model on the dry weight of a female's somatic tissues (somwt).**

| Predictor | | | |
|---|---|---|---|
| Fixed effects: | Estimate (S.E.) | *t*-value | *p*-value |
| Intercept | 4.12 (0.04) | 97.28 | 6.30e-156 |
| Food (high vs. low) | -0.24 (0.03) | -7.45 | 9.05e-12 |
| Predation (high vs. low) | 0.04 (0.04) | 1.10 | 0.27 |
| Dataset (2 vs. 1) | -0.60 (0.05) | -12.98 | 2.31e-25 |
| Drainage (Madamas vs. Mar.) | -0.09 (0.05) | -1.69 | 0.09 |
| Drainage (Madamas vs. Yarra) | -0.18 (0.05) | -3.49 | 0.00065 |
| Random effects: | Var (S.D.) | $\chi_1^2$ | *p*-value |
| Maternal identity | 0.008 (0.088) | 1.08 | 0.30 |
| Residual | 0.066 (0.258) | | |

The dry weight of a female's somatic tissues was measured in milligrams and was base-*e* log-transformed before fitting the model. Model results are provided on the transformed scale. The total number of observations included in the model is 263. As the analysis included data from only four drainages, drainage was fitted as a fixed effect. In dataset 1 fish originated from a single drainage (El Cedro), which was not sampled for any other dataset, so the effects of dataset 1 and the El Cedro drainage cannot be disentangled. S.E.: standard error, Mar: Marianne drainage, Var: variance, S.D.: standard deviation.

**Table S34. Linear mixed-effects model on the female wet weight at the beginning of the experiment (wt0).**

| Predictor | | | |
|---|---|---|---|
| Fixed effects: | Estimate (S.E.) | *t*-value | *p*-value |
| Intercept | 3.52 (0.04) | 87.09 | 5.89e-16 |
| Food (high vs. low) | -0.01 (0.01) | -1.81 | 0.07 |
| Predation (high vs. low) | -0.06 (0.02) | -2.60 | 0.00981 |
| Dataset (2 vs. 1) | 0.30 (0.09) | 3.48 | 0.01749 |
| Dataset (2 vs. 3) | 0.08 (0.04) | 2.21 | 0.02814 |
| Dataset (2 vs. 4) | 0.09 (0.04) | 2.42 | 0.01660 |
| Random effects: | Var (S.D.) | $\chi^2_1$ | *p*-value |
| Maternal identity | 0.034 (0.184) | 305.13 | 2.52e-68 |
| Drainage | 0.005 (0.072) | 14.97 | 0.00011 |
| Residual | 0.011 (0.103) | | |

The female wet weight at the beginning of the experiment was measured in milligrams and was base-*e* log-transformed before fitting the model. Model results are provided on the transformed scale. The total number of observations included in the model is 706. S.E.: standard error, Var: variance, S.D.: standard deviation.

**Table S35. Linear mixed-effects model on the male wet weight at the beginning of the experiment (wt0m).**

| Predictor | | | |
|---|---|---|---|
| Fixed effects: | Estimate (S.E.) | *t*-value | *p*-value |
| Intercept | 3.49 (0.04) | 90.16 | 7.25e-14 |
| Food (high vs. low) | 0.00 (0.01) | 0.49 | 0.63 |
| Predation (high vs. low) | -0.07 (0.03) | -2.74 | 0.00680 |
| Dataset (2 vs. 1) | 0.24 (0.07) | 3.16 | 0.03293 |
| Dataset (2 vs. 3) | 0.13 (0.03) | 3.97 | 0.00013 |
| Random effects: | Var (S.D.) | $\chi^2_1$ | *p*-value |
| Maternal identity | 0.032 (0.178) | 267.75 | 3.52e-60 |
| Drainage | 0.004 (0.060) | 5.78 | 0.01622 |
| Residual | 0.006 (0.078) | | |

The male wet weight at the beginning of the experiment was measured in milligrams and was base-*e* log-transformed before fitting the model. Model results are provided on the transformed scale. The total number of observations included in the model is 458. S.E.: standard error, Var: variance, S.D.: standard deviation.

**Table S36. Linear mixed-effects model on the female wet weight at birth 1 (wt1).**

| Predictor | | | |
|---|---|---|---|
| Fixed effects: | Estimate (S.E.) | *t*-value | *p*-value |
| Intercept | 4.99 (0.05) | 95.94 | 3.43e-13 |
| Food (high vs. low) | -0.28 (0.02) | -15.79 | 9.18e-43 |
| Predation (high vs. low) | 0.08 (0.02) | 3.60 | 0.00037 |
| Dataset (2 vs. 1) | -0.16 (0.12) | -1.35 | 0.24 |
| Dataset (2 vs. 3) | 0.47 (0.04) | 13.26 | 1.25e-31 |
| Dataset (2 vs. 4) | 0.15 (0.04) | 3.88 | 0.00013 |
| Random effects: | Var (S.D.) | $\chi^2_1$ | *p*-value |
| Maternal identity | 0.008 (0.089) | 5.58 | 0.01820 |
| Drainage | 0.011 (0.107) | 50.69 | 1.08e-12 |
| Residual | 0.052 (0.229) | | |

The female wet weight at birth 1 was measured in milligrams and was base-*e* log-transformed before fitting the model. Model results are provided on the transformed scale. The total number of observations included in the model is 686. S.E.: standard error, Var: variance, S.D.: standard deviation.

**Table S37. Linear mixed-effects model on the female wet weight at birth 2 (wt2).**

| Predictor | | | |
|---|---|---|---|
| Fixed effects: | Estimate (S.E.) | *t*-value | *p*-value |
| Intercept | 5.37 (0.05) | 109.56 | 5.72e-13 |
| Food (high vs. low) | -0.28 (0.02) | -17.18 | 4.72e-48 |
| Predation (high vs. low) | 0.06 (0.02) | 2.93 | 0.00360 |
| Dataset (2 vs. 1) | -0.14 (0.12) | -1.24 | 0.27 |
| Dataset (2 vs. 3) | 0.54 (0.03) | 17.17 | 2.20e-46 |
| Dataset (2 vs. 4) | 0.11 (0.03) | 3.23 | 0.00136 |
| Random effects: | Var (S.D.) | $\chi^2_1$ | *p*-value |
| Maternal identity | 0.004 (0.065) | 2.26 | 0.13 |
| Drainage | 0.010 (0.102) | 51.05 | 9.01e-13 |
| Residual | 0.044 (0.210) | | |

The female wet weight at birth 2 was measured in milligrams and was base-*e* log-transformed before fitting the model. Model results are provided on the transformed scale. The total number of observations included in the model is 675. S.E.: standard error, Var: variance, S.D.: standard deviation.

**Table S38. Linear mixed-effects model on the female wet weight at birth 3 (wt3).**

| Predictor | | | |
|---|---|---|---|
| Fixed effects: | Estimate (S.E.) | *t*-value | *p*-value |
| Intercept | 5.69 (0.05) | 122.25 | 5.18e-14 |
| Food (high vs. low) | -0.30 (0.02) | -17.17 | 2.99e-46 |
| Predation (high vs. low) | 0.05 (0.02) | 2.54 | 0.01165 |
| Dataset (2 vs. 3) | 0.54 (0.03) | 17.90 | 1.97e-47 |
| Dataset (2 vs. 4) | 0.00 (0.03) | 0.11 | 0.91 |
| Random effects: | Var (S.D.) | $\chi^2_1$ | *p*-value |
| Maternal identity | 0.002 (0.048) | 0.60 | 0.44 |
| Drainage | 0.009 (0.096) | 51.89 | 5.87e-13 |
| Residual | 0.043 (0.207) | | |

The female wet weight at birth 3 was measured in milligrams and was base-*e* log-transformed before fitting the model. Model results are provided on the transformed scale. The total number of observations included in the model is 583. S.E.: standard error, Var: variance, S.D.: standard deviation.

**Table S39. Linear mixed-effects model on the male wet weight at sexual maturity (wtmat).**

| Predictor | | | |
|---|---|---|---|
| Fixed effects: | Estimate (S.E.) | *t*-value | *p*-value |
| Intercept | 4.11 (0.04) | 100.65 | 4.63e-14 |
| Food (high vs. low) | -0.21 (0.01) | -14.99 | 1.89e-35 |
| Predation (high vs. low) | 0.06 (0.02) | 2.61 | 0.00979 |
| Dataset (2 vs. 1) | 0.02 (0.08) | 0.27 | 0.80 |
| Dataset (2 vs. 3) | 0.40 (0.03) | 13.09 | 1.50e-26 |
| Random effects: | Var (S.D.) | $\chi^2_1$ | *p*-value |
| Maternal identity | 0.017 (0.129) | 45.74 | 1.35e-11 |
| Drainage | 0.005 (0.069) | 18.58 | 1.63e-05 |
| Residual | 0.021 (0.144) | | |

The male wet weight at sexual maturity was measured in milligrams and was base-*e* log-transformed before fitting the model. Model results are provided on the transformed scale. The total number of observations included in the model is 443. S.E.: standard error, Var: variance, S.D.: standard deviation.
